# Non-catalytic role of phosphoinositide 3-kinase in mesenchymal cell migration through non-canonical induction of p85β/AP-2-mediated endocytosis

**DOI:** 10.1101/2022.12.31.522383

**Authors:** Hideaki T. Matsubayashi, Jack Mountain, Tony Yao, Amy F. Peterson, Abhijit Deb Roy, Takanari Inoue

## Abstract

Class IA phosphoinositide 3-kinase (PI3K) galvanizes fundamental cellular processes such as migration, proliferation, and differentiation. To enable multifaceted roles, the catalytic subunit p110 utilizes a multi- domain, regulatory subunit p85 through its inter SH2 domain (iSH2). In cell migration, their product PI(3,4,5)P_3_ generates locomotive activity. While non-catalytic roles are also implicated, underlying mechanisms and its relationship to PI(3,4,5)P_3_ signaling remain elusive. Here, we report that a disordered region of iSH2 contains previously uncharacterized AP-2 binding motifs which can trigger clathrin and dynamin-mediated endocytosis independent of PI3K catalytic activity. The AP-2 binding motif mutants of p85 aberrantly accumulate at focal adhesions and upregulate both velocity and persistency in fibroblast migration. We thus propose the dual functionality of PI3K in the control of cell motility, catalytic and non- catalytic, arising distinctly from juxtaposed regions within iSH2.

## Introduction

Class 1A PI3Ks are lipid kinases that catalyze phosphatidylinositol (3,4,5)-triphosphate (PI(3,4,5)P_3_) production^1, 2^. In the canonical growth factor pathway, PI(3,4,5)P_3_ production leads to Akt/mTOR activation and subsequent upregulation of proliferation and survival. Besides this primary function, PI3K and PI(3,4,5)P_3_ manifest versatile roles in many other physiological contexts including vesicular trafficking, differentiation, immune reaction, and cell migration^2–5^. Due to its multitasking roles, the PI3K catalytic function is modulated by various interaction partners such as ubiquitin ligase Cbl-b^6^, tumor suppressor BRD7^7^, thyroid hormone receptor β^8^, transmembrane tyrosine phosphatase CD148^9^, and microtubule-associated protein MAP4^10^.

Class IA PI3K is a heterodimeric complex composed of a catalytic subunit (p110α, p110β, or p110δ) and a regulatory subunit (p85α, p55α, p50α, p85β, or p55γ)^1, 11, 12^. Upon activation of receptor tyrosine kinases (RTKs), such as platelet-derived growth factor (PDGF) receptors in fibroblasts, nSH2 and cSH2 domains in regulatory subunit recognize tyrosine phosphorylation on the receptors and adaptor molecules^13, 14^. As regulatory subunits tightly associate with p110 through inter SH2 domain (iSH2) that resides between two SH2 domains^11^, p110 consequently accumulates at the plasma membrane. The phosphotyrosine binding of SH2 domains liberates their inhibitory contact with p110^15, 16^, thus resulting in signal-specific PI3K activation proximal to its substrate, phosphoinositide (4,5)-biphosphate.

The catalytic activity of PI3K is one of the major positive regulators in cell migration. In amoeboid cells such as *Dictyostelium discoideum* and mammalian neutrophils, chemoattractant induces PI(3,4,5)P_3_ accumulation at the front of cells^17–19^, leading to the activation of the Rho family of small GTPases including Rac1^19–21^ and cell protrusions driven by the actin cytoskeleton. Mesenchymal cells such as fibroblasts also establish similar PI(3,4,5)P_3_ polarity^22^. However, a recent study found that PI3K in fibroblasts acts as an amplifier of nascent lamellipodia instead of an initiator of protrusion^23^. Further research found that this PI3K-actin feedback loop originates from nascent adhesions, another unique feature of mesenchymal cell migration^24^. Therefore, amoeboid and mesenchymal cells utilize distinct mechanisms, at least at the level of PI3K, with yet elusive mechanisms.

In the face of the catalytic-role-centric studies, non-catalytic roles of p85 have also been reported. In ER stress response, p85 brings XBP-1s to the nucleus to upregulate unfolding protein response genes^25, 26^. p85 also involves in receptor internalization through the interaction with an adaptor molecule insulin receptor substrate 1 (IRS-1), Rab GTPases activation, or ubiquitination on p85 itself^27–29^. In addition, p85 regulates cytoskeletal reorganization in concert with the small GTPase Cdc42^30, 31^. It therefore is important to consider PI3K as a multifaceted molecule to fully understand its functions and regulations. In this study, we combine bioinformatics and chemical biology approaches with live-cell fluorescence imaging to reveal a previously uncharacterized non-catalytic function of PI3K in which a part of the p85β iSH2 domain induces endocytosis mediated by clathrin and dynamin. Using p85 knockout cells with genetic rescues, we show that this non-catalytic induction of endocytosis regulates cell migration properties through local regulation of p85 at focal adhesions.

## Results

### iSH2 domain of regulatory subunit p85 has AP-2 binding motifs

To explore possible non-catalytic roles of PI3K, we analyzed the primary sequence of the regulatory subunits of class IA PI3K (p85α, p85β, and p55γ). Using Eukaryotic Linear Motif (ELM) prediction^32^, we found that iSH2 domain of the C-terminal region of p85β accommodates three consensus binding motifs for AP-2^33^, an adaptor protein for clathrin-mediated endocytosis, namely YxxΦ, di-leucine, and acidic clusters (Fig. 1a, Extended Data Fig. 1). Consistent with the crystal structure of p110 complexed with iSH2-cSH2^16^, the C-terminal region of iSH2 was predicted to be intrinsically disordered and unlikely a part of secondary structures based on primary sequence analysis of IUPred2A^34^, PrDOS^35^, and PONDR^36^ (Extended Data Fig 1). These results suggested possible interaction between p85 and AP-2, which could lead to endocytosis upon their membrane targeting.

**Figure 1:**
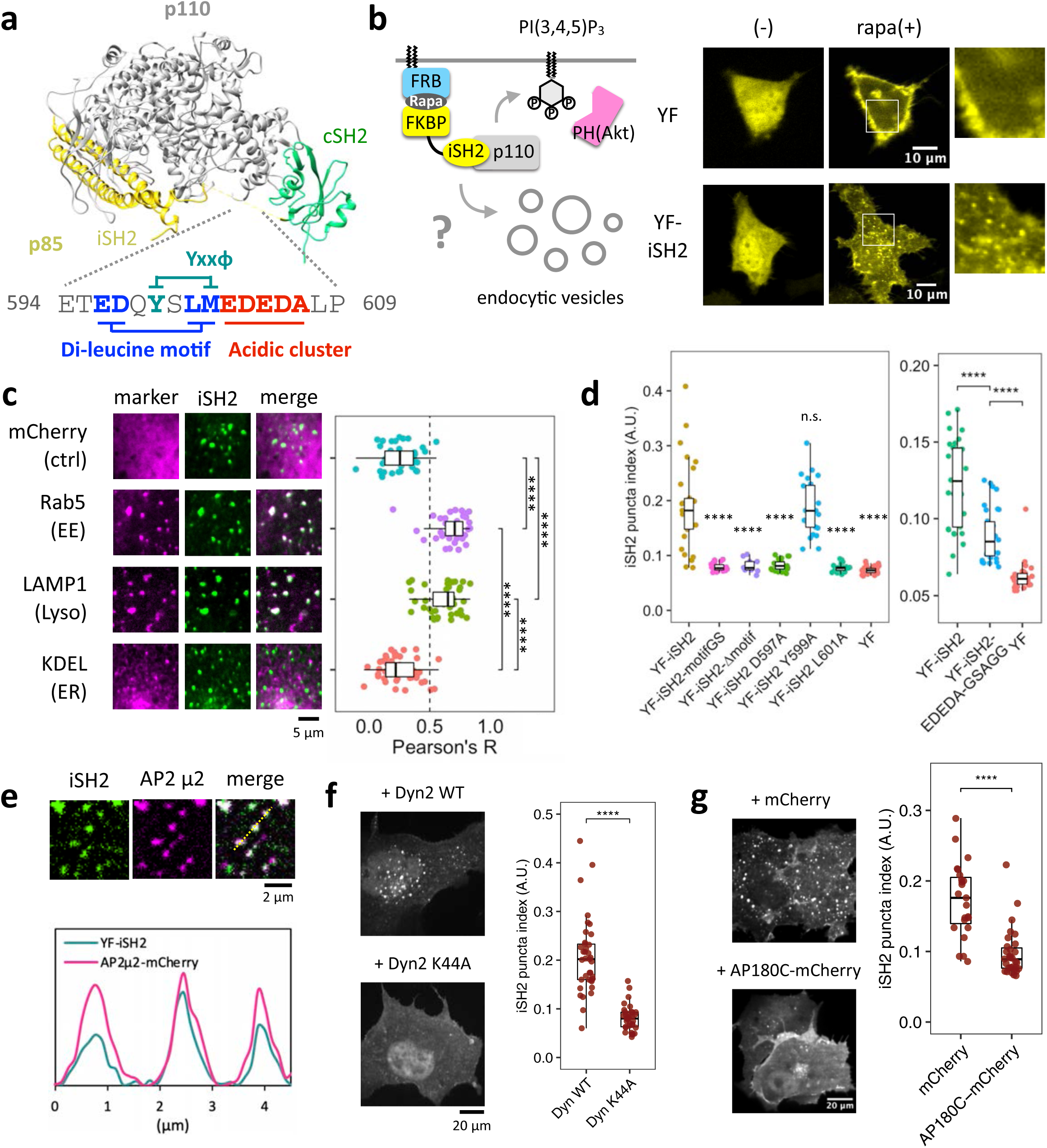
Plasma membrane recruitment of iSH2 domain induces clathrin and dynamin dependent endocytosis. (a) Crystal structure of PI3K (PDB 2y3a) and AP-2 binding motifs of mouse p85β iSH2 domain. (b) Confocal images of endocytic vesicles produced by plasma membrane targeting of iSH2 domain. HeLa cells were transiently transfected with Lyn-ECFP-FRB, mCherry-PH(Akt), and EYFP-FKBP or EYFP-FKBP- iSH2. Images show before and after 100 nM rapamycin addition. (c) Confocal images of iSH2-induced vesicles colocalized with endocytosis marker molecules: mCherry-Rab5 (early endosome) and LAMP1- mRFP (lysosome). mCherry (cytosol) and mCherry-KDEL (ER) were used as negative controls. The graph shows Pearson’s correlation between iSH2 and marker molecules. (d) Quantified iSH2-mediated endocytosis indices (see method) of wild type and mutants in di-leucine motif and acidic cluster, but not YxxΦ motif. (e) TIRF images of iSH2 vesicles colocalized with AP-2. (f, g) Confocal images of iSH2 vesicles showing dynamin and clathrin dependency. Vesicle formation was suppressed in the presence of dominant negative form of dynamin (K44A) or AP180C. Box whisker plots represent median, 1st, 3rd quartiles and 1.5×inter-quartile range. P-values: *: < 0.05, **: < 0.01, ***: < 0.001, ****: < 0.0001. n.s.: not significant. (c, d) Steel-Dwass test. In the right panel of (d), p-values against YF-iSH2 were only shown. (f, g) Wilcoxon rank sum test.

### Plasma membrane recruitment of iSH2 domain induces endocytosis

Whether a given molecule is capable of inducing endocytosis can be tested by recruiting such molecules to plasma membranes^37, 38^. With the help of a chemically inducible dimerization (CID) system^39^, we aimed to recruit iSH2 including the putative AP-2 binding motifs to the plasma membrane and see if this results in endocytosis. To achieve this, we used rapamycin-dependent heterodimerization of FK506-binding protein (FKBP) and FK506-rapamycin-binding domain (FRB) to trap YFP-FKBP-iSH2 (YF-iSH2) at plasma membrane-anchored Lyn-CFP-FRB (Lyn-CR). Within several minutes after accumulation of YF-iSH2 at the plasma membrane, numerous mobile puncta became visible in the cytosol (Fig. 1b, Supplementary movie 1–3). The puncta were seen only with YF-iSH2 but not with a negative control YFP-FKBP (YF), suggesting that iSH2 is responsible for induction of puncta derived from the plasma membrane.

We then tested colocalization between the observed puncta and markers for endocytosis. When we used a membrane staining dye mCLING^40^, which gets internalized to endomembranes upon endocytosis, the puncta colocalized well with the dye (Extended Data Fig. 2). Furthermore, the iSH2 puncta also colocalized with other markers such as mCherry-Rab5 (early endosome) and Lamp1-mRFP (lysosome), but not with negative controls such as mCherry (cytosol) and mCherry-KDEL (ER) (Fig. 1c).

Endocytic activity is highly sensitive to ambient temperature, likely due to critical involvement of dynamin GTPase which has an unusually high Q_10_ temperature coefficient value^41, 42^. When conducting iSH2 recruitment to the plasma membrane at a reduced temperature (37°C to 23°C), we observed much fewer puncta (Extended Data Fig. 3, Supplementary movies 1–3). This is consistent with the lack of documentation of such puncta upon iSH2 recruitment by our group and others in the past^43–46^. Collectively, these results strongly support the idea that membrane-recruited iSH2 induces endocytosis.

### iSH2-mediated endocytosis is context independent

To test how well the iSH2-mediated endocytosis can be generalized, we repeated the CID recruitment assay with two modifications. First, we used FRB anchored to the plasma membrane through six different targeting sequences (Supplementary Table 2). In all cases except KRas4B-CAAX, we observed puncta formation (Extended Data Fig. 4a, b). Furthermore, the endocytosis can be also triggered by a light inducible dimerization system (iLID-SspB)^47^ (Extended Data Fig. 4c). Thus, iSH2-mediated endocytosis is not specific to a certain type of plasma membrane targeting or dimerization scheme.

### iSH2-mediated endocytosis depends on the AP-2 binding motifs

To determine if the predicted AP-2 binding motifs are necessary for iSH2-mediated endocytosis, we deleted 12 amino acids (aa) within the motif clusters (Δmotif) or replaced the same region with a 3×SAGG flexible linker (motifGS). When the recruitment assay was conducted with each of these iSH2 mutants, we saw little to no puncta, indicating the necessity of the 12 aa for inducing endocytosis (Fig. 1d, Extended Data Fig. 5). Then, we individually mutated the YxxΦ motif, di-leucine motif, and acidic cluster. Whereas point mutations in the di-leucine motifs drastically decreased endocytic activity, Y to A mutation in the YxxΦ motif did not show significant effect (Fig. 1d, Extended Data Fig. 5). Replacement of the acidic cluster EDEDA with GSAGG partially reduced the endocytic activity (Fig. 1d, Extended Data Fig. 5). These results suggest that the di-leucine motif and acidic clusters contribute to iSH2-mediated endocytosis.

### iSH2-mediated endocytosis depends on clathrin and dynamin

To understand molecular mechanisms of iSH2-mediated endocytosis, we examined possible association between iSH2 and AP-2 by applying an inducible co-recruitment assay^48, 49^ (Extended Data Fig. 6a). In this assay, we can semi-quantitatively assess a protein-protein interaction in living cells. Here, we recruit an iSH2 domain to the plasma membrane using the chemically inducible dimerization scheme, and measure how much a bait protein, AP-2, gets co-recruited under TIRF microscopy. After recruitment of YFP-FKBP- labelled iSH2 to the plasma membrane, we observed an increase in the fluorescence intensity of AP-2- mCherry (co-recruitment index, CI: 1.23), but not mCherry control construct (CI: 1.03) (Extended Data Fig. 6b, c), implying that iSH2 and AP-2 could interact with each other. This AP-2 co-recruitment was reduced when we used iSH2 motif mutants, Δmotif (CI: 1.07) and motifGS (CI: 1.20) (Extended Data Fig. 6b,c). Similarly, we measured an extent of colocalization between AP-2 and iSH2 after recruitment of iSH2 to the plasma membrane. As a result, AP-2 fluorescence signals on the plasma membrane colocalized with the membrane-recruited iSH2, but not with the motif mutant (Fig. 1e, Extended Data Fig. 6d, e). These results suggested that the AP-2 binding motif of p85 binds to and colocalizes with AP-2 on the plasma membrane.

Interestingly, colocalization of iSH2 and AP-2 was also observed when FRB-CFP-CAAX(KRas4B) was used as a plasma membrane anchor (Extended Data Fig. 6d, e), despite the poor endocytosis induction of CAAX(KRas4B) (Extended Data Fig. 4a, b). This result suggested that while iSH2 interacts with AP-2 regardless of the type of plasma membrane anchor, endocytic development including vesicle maturation and membrane remodeling were somehow stalled in the case of KRas4B-CAAX.

We then tested two dominant negative mutants, N-terminus truncated AP180 (AP180C)^50, 51^ and GTPase- defective dynamin (Dyn2-K44A)^52, 53^, that inhibit endocytic processes. These mutants significantly reduced the numbers of endocytosed puncta, suggesting that iSH2-mediated endocytosis depends on clathrin and dynamin (Fig. 1f, g). Taken together, we conclude that iSH2 brings AP-2 to the plasma membrane, which triggers endocytosis through clathrin and dynamin.

### iSH2-mediated endocytosis is independent of PI3K catalytic activity

Catalytic activity of PI3K and its product PI(3,4,5)P_3_ have been implicated in various types of endocytosis^54–57^. Since the iSH2 domain binds to endogenous p110 and its plasma membrane recruitment leads to PI(3,4,5)_3_ production^43–46^, we asked if iSH2-mediated endocytosis is dependent on PI(3,4,5)P_3_. We tested this with either a PI3K inhibitor (LY294002) or a deletion mutant of iSH2 (iSH2-DN). LY294002 binds to the ATP binding pocket of p110 and inhibit its catalytic function^58^, whereas iSH2-DN mutation abolishes iSH2-p110 interaction^59^. When we performed the iSH2 recruitment assay in the presence of either of these reagents, puncta formation occurred normally despite the production of PI(3,4,5)P_3_ being suppressed in the same cells (Fig. 2a, Extended Data Fig. 7a). This indicates that iSH2- mediated endocytosis is independent of the p110 kinase activity and can be classified as a non-catalytic function of PI3K.

**Figure 2:**
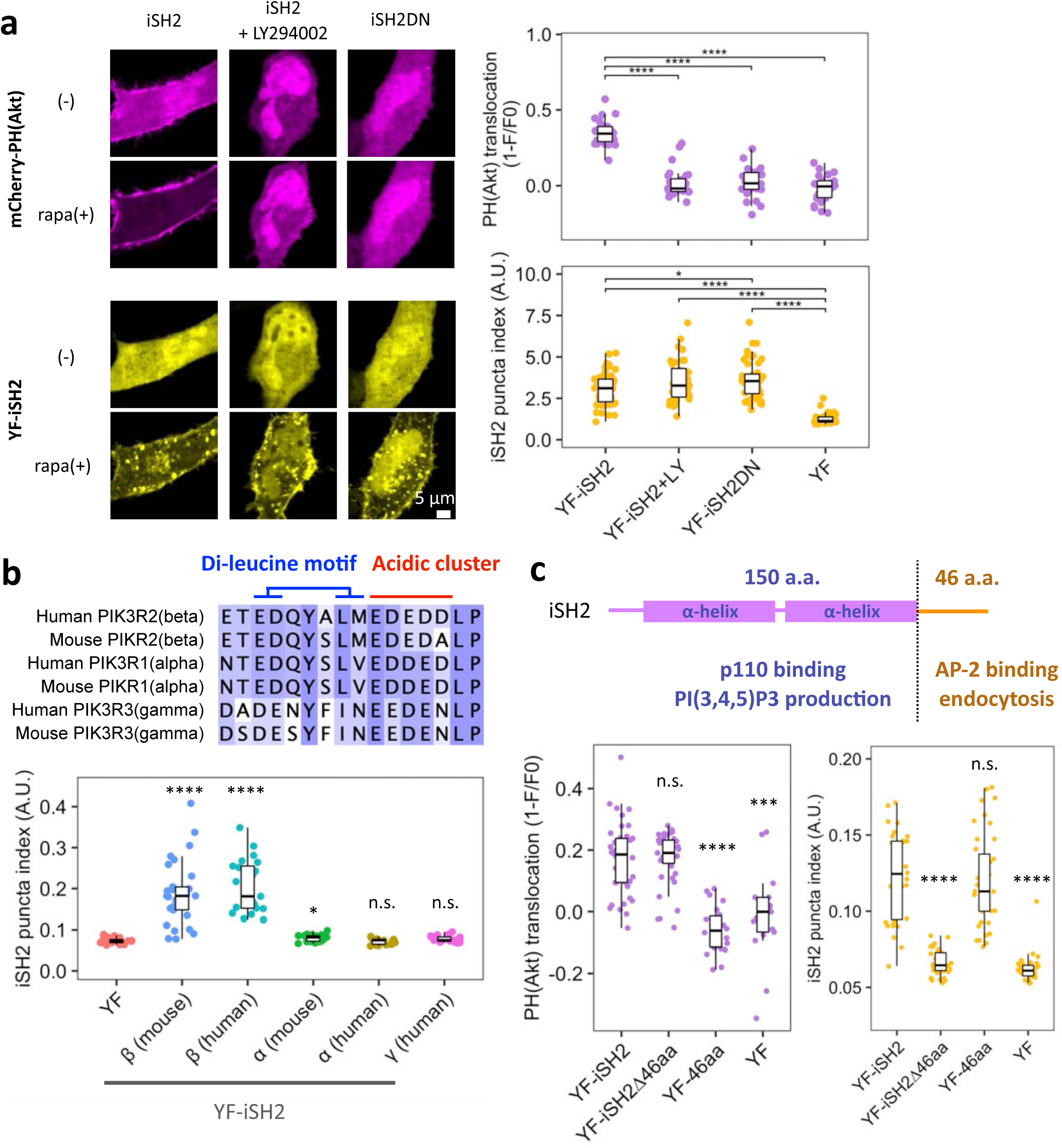
iSH2-mediated endocytosis is independent of PI3K catalytic activity and C-terminal 46 aa region is necessary and sufficient. (a) Confocal images of PI(3,4,5)P3 sensor PH(Akt) and iSH2 vesicles. Quantifications are shown on the right. LY294002: PI3K inhibitor, iSH2(DN): deletion mutant lacking p110 binding site. (b) Top: Amino acid sequence alignment of AP-2 binding motif region of human and mouse p85α, p85β, p55γ isoforms. Bottom: Quantification of iSH2 vesicles produced by each isoform. (c) Secondary structure of mouse p85β iSH2 domain and quantification of PH(Akt) translocation and iSH2 vesicles. Box whisker plots represent median, 1st, 3rd quartiles and 1.5×inter-quartile range. P-values: *: < 0.05, **: < 0.01, ***: < 0.001, ****: < 0.0001. n.s.: not significant.

### iSH2-mediated endocytosis is β isoform specific

The iSH2 domain is defined in all three regulatory subunits of class IA PI3K (p85α, p85β, and p50γ)^1^. We then took iSH2 domains from different isoforms of human and mouse and asked if iSH2-mediated endocytosis is conserved among them by using the CID recruitment assay. iSH2 from p85β (both human and mouse) induced endocytosis, but α or γ isoforms did not (Fig. 2b, Extended Data Fig. 7b), indicating that endocytic activity is β isoform specific. The mechanism of this isoform specificity is unknown, but slight sequential or structural differences may be involved as in the case of the reported isoform-specific binding to Influenza A virus NS1 protein^60–62^.

### 46 aa disordered region is necessary and sufficient for iSH2-mediated endocytosis

The iSH2 domain has been considered as a single domain whose main role is to bind to p110 and bring the catalytic subunit to the plasma membrane upon receptor stimulation. To locate exactly which part of iSH2 contributes to p110 binding, and which part contributes to the endocytosis induction, we performed a sequential truncation to the iSH2 domain. As a result, the C-terminal 46 aa was found to be both necessary and sufficient to induce the endocytosis (Fig. 2c, Extended Data Fig. 7c). In contrast, PI(3,4,5)P_3_ production remained intact with iSH2 lacking this 46 aa region (Fig 2c, Extended Data Fig. 7d, d). Our results demonstrate that the iSH2 domain can be structurally and functionally separated into two regions - the p110 binding coiled-coil region for catalytic actions and the 46 aa disordered region encoding AP-2 motif for non-catalytic induction of endocytosis.

### Generation of MEF cell lines with p85β AP-2 binding motif mutants and their biochemical characterization

To investigate how the unexpected link between p85β and AP-2 influences the cellular functions of PI3K, we took an advantage of p85α/β double knock out (DKO) in mouse embryonic fibroblasts (MEFs)^63^ to which a series of p85 variants, with or without mutations in AP-2 binding motifs, were individually introduced via lentiviral infection (Extended Data Fig. 8a). Since both the di-leucine motif and the acidic cluster contribute to endocytic activity (Fig. 1d), we created two p85β mutants whose 12 aa motif region was either truncated or replaced with 3×SAGG, serving as AP-2 motif deficient forms of p85β. YFP was tagged on the rescued p85 to sort the virus-infected cells and validated the consistency in the expression level of rescued p85 variants (Extended Data Fig. 8b).

Using these genetic resources, we first assessed a possible regulatory role of the AP-2 binding motif in a receptor tyrosine kinase pathway (Fig. 3a). Consistent with a previous report^63^, expression of wild type p85β in DKO MEFs could rescue the elevated levels of Akt phosphorylation (pTyr-308) in response to PDGF addition (Fig 3b). When we tested this with the mutant p85β cell lines, there was no significant difference from the wild type. In assessing cell proliferation, we then found similar proliferation rates for cells rescued with wild type and motifGS mutant (Fig. 3c). Thus, mutations in the AP-2 binding motif of p85β did not show an apparent effect on Akt response or cell growth. Considering the possibility that AP- 2 binding of p85β regulates receptor internalization, we next measured the effect on ERK, the other major pathway regulated by endocytic traffic of receptor tyrosine kinase (RTK)^64^. However, wild type and mutant rescued cells showed a similar pattern in ERK response (Extended Data Fig. 8c). We also tested the effect on transferrin receptors, a typical cargo of clathrin-dynamin endocytosis, and found no significant change in transferrin internalization between wild type and mutant rescue cells (Extended Data Fig. 8d). Therefore, the binding between p85β and AP-2 did not seem to influence on RTK signaling or general endocytic functions.

**Figure 3:**
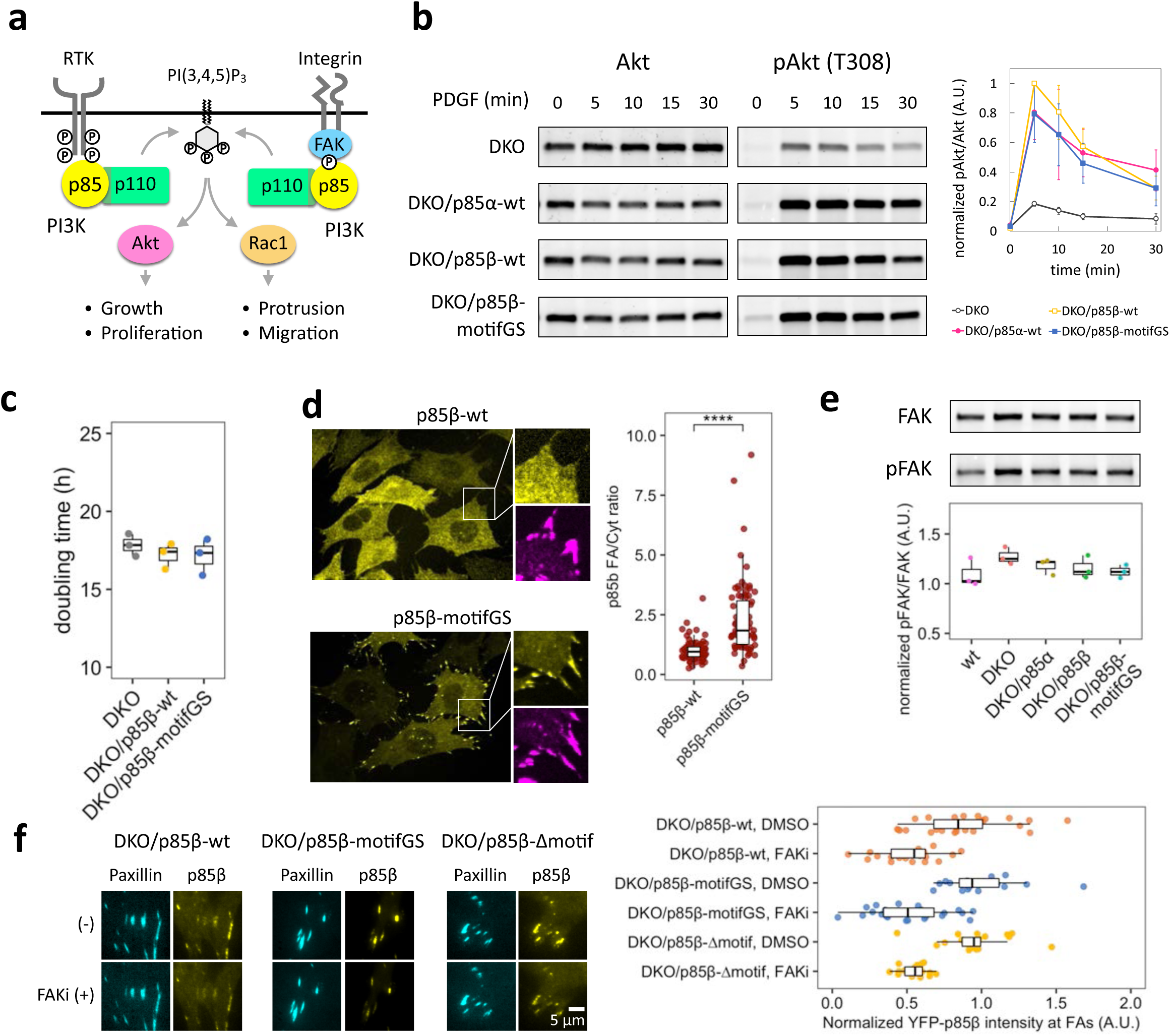
Mutation in AP-2 binding motifs of p85β increases focal adhesion localization. (a) Schematic of receptor tyrosine kinase-dependent and focal adhesion-dependent PI3K pathways. (b) Western blot of total- and phospho-Akt (T308) and its quantification. Cells were treated with 50 ng/mL PDGF for indicated time. pAkt/Akt level was normalized to DKO/p85β-wt 5 min. Error bars represent standard deviations. (c) Doubling time of DKO and p85 rescued MEF cells. (d) Confocal images of p85β-wt and p85β-motifGS cells and their quantification. Yellow: EYFP-p85β, Magenta: immunofluorescence against vinculin. (e) Western blot of total- and phospho-FAK (Y397) and its quantification. (f) FAK activity dependency of p85 focal adhesion localization. Cells were treated with DMSO or 10 µM PF-573228 (FAK inhibitor; FAKi) for 5 min and EYFP-p85β intensity were divided by the values of time=0. Box whisker plots represent median, 1st, 3rd quartiles and 1.5×inter-quartile range. P-value:****: < 0.0001. (d) Wilcoxon rank sum test.

### Mutations in AP-2 binding motif causes localization of p85β at focal adhesions

Besides the RTK response, PI3K locally controls cellular morphodynamics in association with focal adhesions^24, 30, 65, 66^. To determine if AP-2 binding motifs are involved in such subcellular regulation, we next investigated the intracellular localization of wild type and mutant p85β using confocal microscopy. Strikingly, the 3×SAGG and Δmotif p85 cell lines showed significantly enhanced accumulation at focal adhesions (Fig. 3d). Previous studies found that p85 localizes to focal adhesions where it binds to focal adhesion kinase (FAK) through the interaction between its SH3 domain and auto-phosphorylated tyrosine of FAK (pY397)^65, 67–70^. We thus tested the effect of the AP-2 motif mutation on FAK. Western blot analysis did not detect significant differences in the expression or phosphorylation level of FAK among the p85-rescued cell lines (Fig. 3e). Using TIRF microscopy, we further performed live-cell imaging of p85 fused to YFP which was co-expressed with a focal adhesion marker mCerulean3-Paxillin^71^ in the presence or absence of an FAK inhibitor PF-573228^72^. The results showed that both wild type and mutant p85 dissociated from focal adhesions after FAK inhibition with identical kinetics (Fig. 3f, Extended Data 9). Together, the data suggest that AP-2 binding motifs are involved in sequestration of p85β from focal adhesions. Since the observed sequestration did not affect the interaction between the SH3 domain of p85β and pY397 of FAK, there is another mechanism underlying a trigger of the sequestration.

### Fibroblasts with impaired AP-2 binding motifs migrate faster and more persistently

Focal adhesions function as a molecular clutch for a cell to transmit mechanical force to the external environment^73^, while simultaneously serving as a biochemical hub for PI3K-Rho GTPase-actin to extend lamellipodial protrusion^24, 66^. Since mutation in AP-2 binding motifs altered localization of p85β at focal adhesions, we hypothesized that AP-2 binding motifs regulate cell migration through focal adhesions. To test this, we characterized migratory properties in a series of DKO MEFs in the presence of 10% FBS to trigger random migration (Fig. 4a, Extended Data Fig. 10a). DKO MEFs exhibited slower migration speed than wild type counterpart MEFs (Fig. 4b), consistent with the reduced Rac activity and less lamellipodia formation in the knockout cells^63^. Interestingly, rescuing the DKO cell line with wild type p85β further decreased migration speed (Fig. 4b, c). In contrast, the cells rescued with AP-2 binding motif mutants of p85β or p85α did not show the decrement, suggesting that the AP-2 motif negatively regulates migration (Fig. 4b, c, Extended Data Fig. 10).

**Figure 4:**
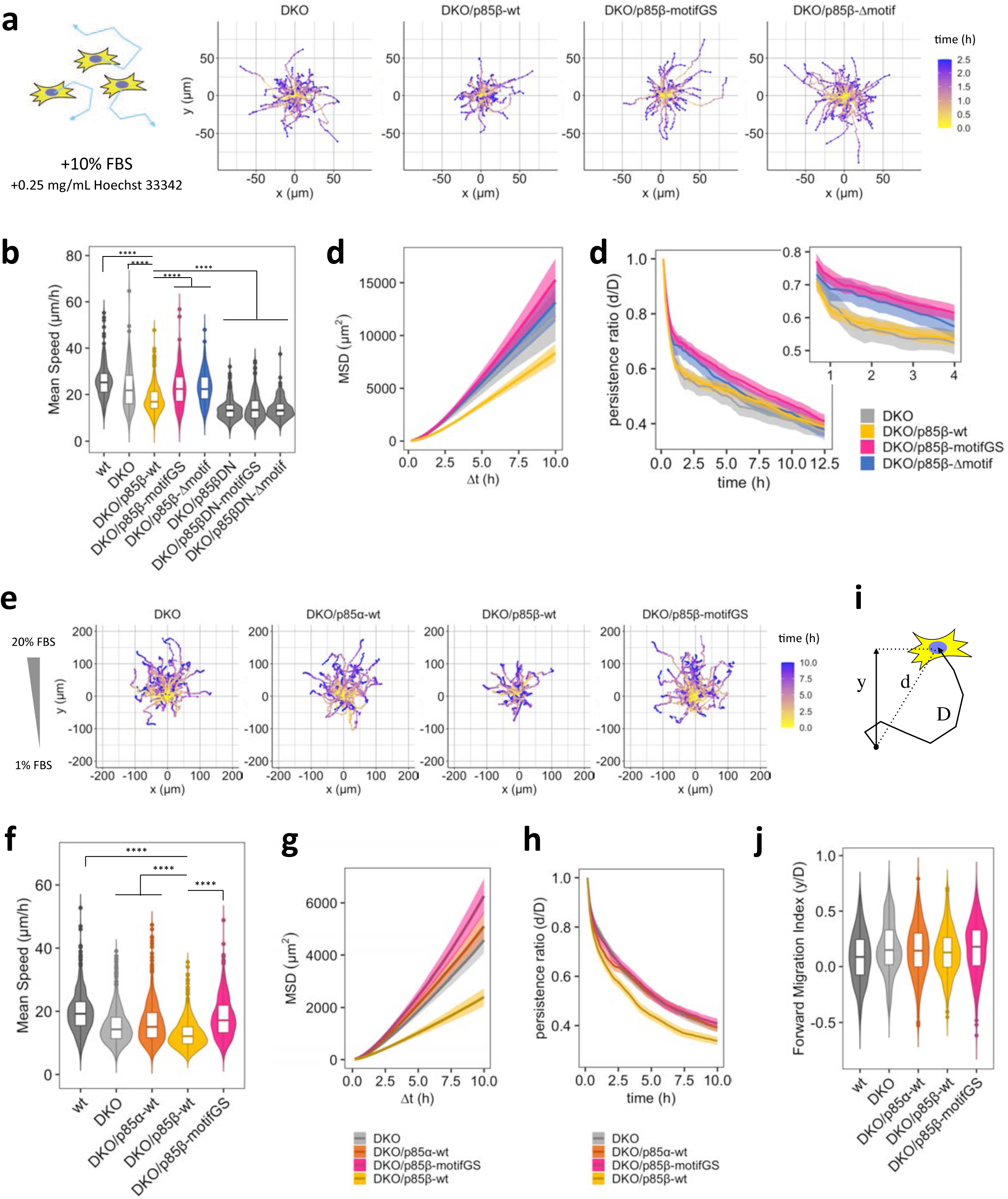
Mutation in AP-2 binding motifs of p85β enhances cell motility in random and chemotactic migration. (a) Representative tracks of 2D random migration on fibronectin coated plates. Cells were allowed to migrate at 37°C with 5% CO_2_ and 10% FBS. 0.25 mg/mL Hoechst 33342 was used for tracking cells. (b, c, d) Quantification of migration parameters. Error bars in (c) and (d) represent 2×SEM (95% CI). (e) Representative tracks of chemotaxis in µ-Slide chemotaxis chamber (ibidi). Cells were allowed to migrate at 37°C with 5% CO_2_ in the presence of 1–20% FBS gradient. 0.25 mg/mL Hoechst 33342 was used for tracking cells. (f, g, i and j) Quantification of migration parameters. Error bars in (g and i) represent 2×SEM (95% CI). (h) Schematic of displacement: d, distance: D, and forward displacement: y. Persistence ratio was defined as d/D, while Forward migration index was defined as y/D. Box whisker plots represent median, 1st, 3rd quartiles and 1.5×inter-quartile range. (b, f, and j) Steel-Dwass test was performed and p-values against DKO/p85β-wt were indicated. P-values: ****: < 0.0001. n.s.: not significant. In (j), p-values of Steel-Dwass test were < 0.001 for wt-DKO, <0.05 for wt-DKO/p85α-wt, a <0.001 for wt-DKO/p85β-motifGS, respectively, while the other pairs were not significant.

Dominant negative mutation of p85 (DN), which lacks 470 to 504 aa residues necessary for p110 binding and decouples catalytic activity of PI3K from receptor activation^59^, and pharmacological inhibition of PI3K and FAK completely suppressed the migration. This basal level of migration was significantly lower than the migration activity of wild type p85β-rescued cells (Fig. 4b, Extended Data Fig. 10a, b). These results suggest that p85β has two layers of regulations on cell migration: positive regulation through PI3K catalytic product, PI(3,4,5)P_3_ and negative regulation through AP-2-mediated sequestration of p85β from focal adhesions.

We then calculated persistence ratio of cell motility defined as the ratio between displacement (d) and the total path length (D), which decreased over the course of migration assays. The decrease in wild type p85β-rescued cells was more prominent over time than mutant p85-rescued cells, suggesting that the link between p85 and AP-2 is involved in a negative regulation of cell migration with a temporal delay from PI(3,4,5)P_3_-mediated positive regulation (Fig. 4d, Extended Data Fig. 10c). Difference in migration speed between wild-type p85 rescue cells and AP-2 motif mutant rescue cells was also seen with PDGF as a stimulant, instead of FBS (Extended Data Fig. 10d), suggesting that the AP-2-mediated motility control is at play under growth factor signaling.

### Role of the AP-2 binding motif in chemotaxis

To test migration behavior in a physiologically relevant context, we performed chemotaxis assays where cells are guided to migrate in a directed manner according to a chemoattractant gradient (Fig. 4e). In line with the random migration results, p85β-rescued cells migrated more slowly than that of DKO, p85α- rescued, and p85β motif mutant-rescued cells (Fig. 4f, g). Although the persistent ratio drew slightly different curves from those of random migration, wild type p85β-rescued cells consistently showed the least persistency among the tested cells (Fig. 4h). These data support the negative regulation of chemotaxis by the AP-2-mediated endocytosis. To examine its role in gradient sensing during chemotaxis, we quantified the forward migration index (FMI) defined as a ratio between forward displacement (y) and the total path length (D) (Fig. 4i). As a result, there was no significant difference in FMI among the conditions tested; wild type cells, DKO cells, and DKO cells rescued with p85α, p85β, or p85β-motifGS (Fig. 4j). These data suggest that the AP-2-mediated endocytosis downregulates migration properties such as speed and persistence, but not gradient sensing, during chemotaxis.

## Discussion

The iSH2 domain is characterized as a positive regulator of PI3K since it stabilizes and recruits the catalytic subunit p110 to the plasma membrane^74^. Our present study demonstrates that the iSH2 domain of p85β has concurrent negative regulation of cell migration through AP-2-mediated endocytosis which originates from the C-terminal disordered region. Disruption of this linkage between p85β and AP-2 led to abnormal accumulation of p85β at focal adhesions (Fig. 3) and also increased speed and persistency of cell migration (Fig. 4). Based on these findings, we propose that the iSH2 domain, originally assigned as a single domain for a single function, consists of two parts with distinct, antagonistic functions: the p110 binding coiled-coil region to promote cell migration, and the AP-2 motif-encoding disordered region to induce endocytosis for negative regulation of cell migration. One may wonder why PI3K elicits two opposing signals for cell motility control. Such a seemingly meaningless regulation may be explained by the kinetic difference. Upon stimulation, PI(3,4,5)P_3_ production can initiate within milli-seconds to seconds timescale^75^, while clathrin-mediated endocytosis occurs more gradually (tens of seconds to a few minutes)^76^. The temporal difference creates an autonomous delayed negative feedback loop, which is one of the signature characteristics necessary for self-organized signal transduction often proposed in directed cell migration^77^. Thus, for PI3K to send out counteracting signals of different kinetics may be of importance for this intricate cell function.

We also determined that AP-2 motif regulates p85β localization at focal adhesions. Since cell protrusion signaling consisting of PI3K and actin is closely coupled with cell adhesions^23, 24, 66^, sequestration of PI3K from focal adhesions could act as a negative regulator of chemotaxis. Considering that mutations to the AP-2 binding motif did not affect the expression level or FAK phosphorylation (Fig. 3), the p85-mediated endocytosis likely regulates the signals downstream of PI3K without drastically altering molecular composition of the focal adhesions. Interestingly, under PDGF stimulation, mutations in the AP-2 binding motif increased cell migration speed without affecting other major pathway effectors such as Akt and ERK (Fig. 3b, c, Extended Data Fig. 8c, 10d). How does AP-2-mediated regulation discriminate a specific signaling molecule from others? Two interesting observations may be of help to answer this question - the AP-2 binding motif resides within the intrinsically disordered region (Extended Data Fig. 1), and many of the membrane anchors that led to the p85-mediated endocytosis (Extended Data Fig. 4) colocalize with ordered lipid domains. Both properties are known to form unique molecular organizations such as liquid droplets and lipid rafts. It is thus intriguing to speculate that it is this unique lipid-protein interaction that results in biomolecular organization prerequisite for the p85-mediated endocytosis.

PI3K activity at focal adhesion is a major driver of mesenchymal cell migration. Earlier works showed that mesenchymal cells initiate protrusion with filopodia extension from nascent adhesions and that a positive feedback loop consisting of PI3K and actin dilates these adhesion-associated protrusions to develop mature lamellipodia^23, 24^. Given that p85β has greater affinity to focal adhesion than p85α^65^, p85β is assumed to play a dominant role in cell migration. We determined that AP-2 binding of p85β negatively regulates its focal adhesion residence. As extension/retraction of membrane protrusions and their lifetime are all proportional to the PI3K activity^78^, this AP-2-mediated sequestration of p85β could act as a brake for migrating cells. Indeed, our data indicated that speed and persistency of cell migration correlate with extent of p85β localization at focal adhesions. Furthermore, the AP-2-mediated sequestration could fulfill a condition for long-sought negative feedback regulation of the PI(3,4,5)P_3_ excitability^24^. Further exploration of molecular mechanisms underlying the observed p85β dissociation from focal adhesion should help reveal the understudied negative feedback regulation.

Of great interest, iSH2-mediated endocytosis is specific to the β isoform and not observed with α or γ isoforms. Their opposing effects are reported elsewhere. For instance, p85α and p85β act as a tumor-suppressor and an oncogene, respectively^65, 79–82^. Such a difference may have something to do with the endosomal PI3K signaling driven by p85β, but not by p85α. Recent studies revealed a role of endosomal PI(3,4,5)P_3_ in Akt signaling^10, 83^. In addition, the AP-2 binding motif region coincides with the hinge region that determines the oncogenicity of p85β^82^. Thus, iSH2-mediated endocytosis possibly contributes to hyperactivate endosomal PI3K-Akt signal. T cell regulation may also be a target of p85β endocytosis. It was shown that T cell coreceptor CD28 preferentially binds to the p85β isoform^84^, and that a PI3K- dependent endocytic process determines the CD28 pathway activity^85^. It is therefore tempting to speculate that iSH2-mediated endocytosis associates with the enigmatic difference in immune phenotypes between p85α and p85β knockout mice^1, 11, 86, 87^. Accordingly, the impact of p85β-mediated endocytosis on physiological functions, as well as the molecular mechanisms leading to the difference between α and β, are fundamental to comprehensive understanding of the multi-faceted PI3K molecule in both normal and cancer cells.

## Author Contributions

HTM initiated the project. HTM, JM, ADR and TI designed the experiments. HTM, JM, TY, AP and ADR performed the experiments and analyzed the data under the supervision of TI. HTM wrote the manuscript in consultation with TI. HTM, JM, and TI edited the manuscript. All the authors contributed to the final version of the manuscript.

## Acknowledgement

We thank Brendan Manning for p85 double knockout cells; Andrew Ewald for HEK293FT cells; Sandra B. Gabelli for human p85α plasmid; Gerald R.V. Hammond for Rab5 and LAMP1 plasmids; Justin W. Taraska for AP180 plasmid; Yi Wu for Paxillin plasmid. We also thank Yuta Nihongaki for technical assistance on lentivirus and FACS experiments. We appreciate Yoshihiro Adachi, Hiroshi Senoo, Miho Iiijma, and Hiromi Sesaki for technical support on lentivirus and western blot experiments. Our appreciation extends to Shigeki Watanabe, Yuuta Imoto, Atsuo Sasaki, Sho W. Suzuki, and Chuan-Hsiang Huang for insightful comments on the research project, and to our lab members including Hideki Nakamura, Allister Suarez and Helen D. Wu for fruitful discussions. We also thank Robert DeRose for manuscript proofreading and experimental support. This study was supported by the National Institutes for Health (R01GM123130 and R01GM136858 to TI, T32GM007445 to AFP), the DoD DARPA (HR0011-16-C-0139 to TI), and the PRESTO program of the Japan Science and Technology Agency to HTM (JPMJPR20KA). HTM was supported by Postdoctoral Fellowships from the Japan Society for the Promotion of Science.

## Materials and Methods

### Reagents and antibodies

Rapamycin was purchased from LCLab (R-5000), prepared as 100 µM stock solution in DMSO, and stored at -20°C. Alexa Fluor 647 conjugated transferrin was purchased from Thermo Fisher Scientific (T23366), reconstituted with Milli-Q water to obtain 5 mg/mL stock solution in PBS, and stored at 4°C. mCLING-ATTO 647N-labeled was purchased from Synaptic Systems (710 006AT1), reconstituted with Milli-Q water to obtain 50 µM stock solution in PBS, and stored at -80°C. LY294002 was purchased from Selleck Chemicals (S1105), prepared as 50 mM stock solution in DMSO, and stored at -20°C. Fibronectin was purchased from Sigma-Aldrich (F4759), reconstituted with Milli-Q water to obtain 1 mg/mL stock solution, and stored at -20°C. Once frozen fibronectin was thawed, the remainder was kept at 4°C. PDGF-BB was purchased from Sigma-Aldrich (P3201), reconstituted with 4 mM HCl containing 0.1% BSA to obtain 50 µg/mL stock solution, and stored at -20°C. FAK inhibitor PF-573228 was purchased from Selleck Chemicals (S2013), prepared as 20 mM stock in DMSO, and stored at -20°C. Hoechst 33342 (10 mg/mL solution in water) was purchased from Thermo Fisher Scientific (H3570) and stored at 4°C. Vinculin antibody (MAB3574-25UG) was purchased from Sigma-Aldrich. Akt (9272S), phospho-Akt (T308) (13038S), FAK (13009S), and phospho-FAK (Y397) (8556S) antibodies were purchased from Cell signaling. GAPDH antibody (sc-32233) was purchased from Santa Cruz. Alexa Fluor 488-conjugated anti-Rabbit IgG (A- 21206), Alexa Fluor 568-conjugated anti-Mouse IgG (A11004), Alexa Fluor 647-conjugated anti-Mouse IgG (A-31571), and Alexa Fluor 647-conjugated transferrin (T23366) were purchased from Thermo Fisher Scientific.

### Plasmids

The sequence of Lyn^88^, KRasCAAX^89^, EYFP-FKBP^90^, EYFP-FKBP-iSH2β(mouse), and PH(Akt)^43^ have been reported elsewhere and their plasmids are summarized in Supplementary Table 1. The other plasma membrane anchors were constructed based on Lyn-ECFP-FRB or FRB-ECFP-KRasCAAX by replacing membrane anchor sequences with synthesized oligo DNA. ORF sequences of the plasma membrane anchor series are summarized in Supplementary Table 2^91–93^. Of note, LAT-ECFP-FRB was tagged with Kir2.1 signal (RAQLLKSRITSEGEYIPLDQIDINVGFDSG) and ER export signal (NANSFCYENEVALTSK) to maximize plasma membrane localization^94^. EYFP-FKBP-iSH2β(mouse)-DN was constructed by deleting M470–R504 by inverse PCR with the primer set (fwd: 5’-GCTGCAGCGAGAGGGAAATGAGAAG-3’, rev: 5’- CCTCTCGCTGCAGCTCCTGGGAGGT-3’). iSH2β(mouse)-Δmotif was PCR-amplified with template plasmid EYFP-FKBP-iSH2β(mouse) and the primer set (fwd:5’- GCTGGTGGTCCTCGAGCATCCAAGTACCAACAAGACCAGG-3’, rev: 5’-AATTGAATTCTCAAGTCTCGTTCTTGATTCCCAG-3’) and inserted between XhoI and EcoRI sites by restriction digestion and T4 ligation. iSH2β(mouse)-motif-3×SAGG was similarly PCR-amplified with the template plasmid EYFP-FKBP-iSH2β(mouse) and with the primer set (fwd:5’- GCTGGTGGTCCTCGAGCATCCAAGTACCAACAAGACCAGG-3’, rev: AATTGAATTCTCACGTGCGCTCCTCGTGGTGGGGGAGGCCTCCGGCAGACCCGCCTGCGGAGCCTCCAGCGCTAGTCTCGTTCTTGATTCCCAG) and inserted between XhoI and EcoRI sites. Alanine mutants of motif sequences were created by inverse PCR with corresponding primer sets.

mCherry-Rab5(*C. lupus*) and LAMP1(human)-mRFP were kind gifts from Dr. Gerald R.V. Hammond. mCherry-KDEL, mCherry-Dyn(WT), and mCherry-Dyn(K44A) were constructed by replacing the fluorescent protein part of YFP-KDEL^95^, YFP-Dyn(WT), and YFP-Dyn(K44A)^89^ with restriction digestion and T4 ligation. AP180(rat)-mCherry was a kind gift from Dr. Justin W. Taraska. To make the truncated version AP180C-mCherry, AP180 (530–918 aa) was PCR-amplified with the primer set (fwd: 5’- CTTCGAATTCTGGCCACCATGGCTGCCGCCACCACC-3’, rev: 5’- CGGTGGATCCccCAAGAAATCCTTGATGTTAAGATCCGCTAATGG-3’) and inserted into EcoRI and BamHI sites of pmCherry-N1 (Clontech) by restriction digestion and T4 ligation. AP2µ2(rat)-mCherry was obtained from Addgene (#27672).

The plasmids of mouse p85α, human p85β, and human p55γ were obtained from Addgene (#1407, #70458, # 70459). The plasmid of human p85α was obtained from DNASU. To construct EYFP-FKBP- iSH2α(mouse), EYFP-FKBP-iSH2α(human), EYFP-FKBP-iSH2β(human), and EYFP-FKBP-iSH2γ(human), each iSH2 region was PCR-amplified with the template of corresponding p85 or p55 plasmid and the primer sets (mouse-α-fwd: 5’-GGTCCTCGAGCATCCAAATACCAGCAGGATCAAGTTG-3’, mouse-α-rev: 5’-GTCGACTGCAGAATTCTCAGGTTTTCTCATCATAATGGGGC-3’) and inserted between XhoI and EcoRI sites of EYFP-FKBP by restriction digestion and T4 ligation or Gibson assembly. EYFP-p85β(mouse) was constructed by inserting PCR-amplified p85β(mouse) (fwd: 5’-AGATCTCGAGCTAGTGCTGGTGGTAGTGCTGGTGGTAGTGCTGGTGGTAGTGCTGGTGGTAGTGCTGGTGGTATGGCAGGAGCCGAGG-3’, rev: 5’-TGCAGAATTCTCAGCGTGCTGCAGACG-3’) between XhoI and EcoRI with restriction digestion and T4 ligation. EYFP-p85β(mouse)-motifGS was consttucted by inverse PCR and T4 ligation with the primer set pretreated with T4 polynucleotide kinase (fwd: 5’- GGCGGGTCTGCCGGAGGCCTCCCCCACCACGAGGA-3’, rev: 5’- TGCGGAGCCTCCAGCGCTAGTCTCGTTCTTGATTCCCAGC-3’). EYFP-p85β(mouse)-Δmotif, EYFP-p85β(mouse)-DN (deletion of M470–R504) were created by inverse PCR with the primer sets (motifGS- fwd:, motifGS-rev:, Δmotif-fwd: 5’-ACGAGACTCTCCCCCACCACGAGGAG-3’, Δmotif-rev: 5’- GGGGGAGAGTCTCGTTCTTGATTCC-3’, DN-fwd: 5’-GCTGCAGCGAGAGGGAAATGAGAAG-3’, DN-rev: 5’-CCTCTCGCTGCAGCTCCTGGGAGGT-3’). For lentivirus vector construction, EYFP-p85 and its mutants were subcloned into FUGW-puro lentivector (a kind gift from Reddy lab) by using AgeI and EcoRI sites. To construct FUGW-puro-Paxillin(human)-mCerulean3, human Paxillin sequence was PCR-amplified from the template pTriEx-mCherry-Paxillin (a kind gift from Yi Wu lab) with the primer set (fwd: 5’- ATCCCCGGGTACCGGGCTAGCGCCACCATGGACGACCTCGACGCCC-3’, rev: 5’- CATGGTGGCGACCGGTGAACCAGCACTACCACCAGCACTACCACCAGCACTACCACCAGCACTGCAGAAGAGCTTGAGGAAGCAG-3’) and inserted into AgeI site of FUGW-puro lentivector by Gibson assembly.

### Cell culture

HeLa, Cos-7 and HEK293FT cells (a kind gift from Andrew Ewald lab) were cultured in a DMEM (Corning, 10-013-CV) medium supplemented with 10% fetal bovine serum (Sigma-Aldrich, F6178). Wild type and p85 double knock out (DKO) mouse embryonic fibroblast (MEF) cells were kind gifts from Brendan Manning lab and cultured in DMEM with 10% FBS.

### Generation of YFP-p85 rescued MEF cells

EYFP-p85 rescued cells were established by lentivirus transduction. Lentiviruses were produced by transfecting HEK293FT cells as follows. Five hundred micro litter of Opti-MEM was mixed with 10 µg FUGW-puro-EYFP-p85, 7.5 µg Δ8.9, and 3.5 µg VSV-G plasmids. Another 500 µL of Opti-MEM was mixed with 63 µL of 1 mg/mL polyethylenimine. Two solutions were mixed and kept at room temperature for 20 minutes, then added to HEK293FT cells seeded one day before at 6×10^6^ cells/10 cm dish density. Two and three days after transfection, media were collected. The virus-containing media were mixed with 1/3 volume of 40% (w/v) PEG-8000, 1.2 M NaCl, 1×PBS (pH 7.0–7.2) and kept at 4°C for more than 45 min. The viruses were precipitated by centrifugation (1,500×g for 45 min at 4°C) and resuspended with PBS (200 µL for 10 cm dish cells). Aliquoted viruses were flash-frozen in liquid nitrogen and stored at -80°C. To infect p85 DKO cells with the viruses, p85 DKO cells were seeded one day before infection at 4×10^4^ cells/well (6-well) density. On the day of infection, medium was replaced with fresh 500 µL of medium and virus suspension (10–100 µL depending on titer) and final 10 µg/mL polybrene were added. YFP positive cells were sorted by SH800S (SONY).

### Transient transfection

HeLa and Cos7 cells were transfected by lipofection with XtremeGene9 (Sigma-Aldrich, 6365787001) in reverse transfection manner. Typically, 40 µL Opti-MEM, 1 µL XtremeGene9, and 0.5–1 µg of plasmid DNA were used for 2 wells (8-well, 75×10^3^ cells/well for Cos7 cells, 150–200×10^3^ cells/well for HeLa cells, 25– 50×10^3^ cells/well for MEF cells) and incubated at 37°C with 5% CO2 and 95% humidity, for 16–24 hours before imaging. 8-well chambers (154534) were poly-D-lysine (P6407-5MG) coated except for TIRF AP-2 colocalization assay (strong adhesion stabilizes AP-2 on the plasma membrane and interferes with the imaging). MEF cells were transfected either by lipofection with XtremeGene9 or by electroporation with Nucleofactor 2b. For electroporation, 2×10^6^ cells were resuspended with Nucleofactor kit T solution (+ supplement 1) and mix with 5 µg plasmid DNA. After zapping with T-20 protocol, 1 mL culture medium was quickly added to the samples and the cells were seeded on fibronectin coated 8-well chambers at the density of 25–50×10^3^ cells/well.

### Microscopes and imaging

Confocal imaging was performed on a spinning-disk confocal microscope. The microscope was based on an inverted Axiovert 200 microscope (Zeiss) and equipped with the spinning disk confocal unit (CSU10; Yokogawa) and triple-band dichroic mirror (Di01-T442/514/647, Semrock). Excitations of CFP, YFP, and mCherry were conducted with diode lasers and a semiconductor laser (COHERENT, OBIS 445 nm LX 75 mW, OBIS 514 nm LX 40 mW, OBIS 561 nm LS 50 mW), which were fiber-coupled (OZ optics) to the spinning disk unit. Images were taken with a Neo Fluor ×40 objective (Zeiss) and a CCD camera (Orca ER, Hamamatsu Photonics) driven by or MetaMorph or Micro-Manager 1.4 (Open Imaging). Images of live cell CID assay was typically taken every 1 min for 40 min. Epi imaging for mCLING assay sample and ERKKTR live cell Imaging was performed by an Eclipse Ti inverted fluorescence microscope (Nikon) equipped with a ×60 oil-immersion objective lens and Zyla 4.2 plus sCMOS camera. TIRF imaging of focal adhesion was performed by an Eclipse Ti inverted fluorescence microscope (Nikon) equipped with a ×100 oil-immersion TIRF objective lens and pco.edge sCMOS camera (PCO). Nikon microscopes were driven by NIS-Elements software (Nikon).

All the live cell imaging was performed in the imaging media containing DMEM (Corning, 17-205-CV) and 1×Glutamax (Thermo Fisher Scientific, 35050061) with temperature (37°C), CO2 (5%), and humidity control by a stage top incubator and a lens heater (Tokai Hit). For fixation, typically, cells were chilled on ice, washed 2 times with ice-cold PBS, fixed by fixation solution (4% paraformaldehyde and 0.15 % glutaraldehyde in PBS) for 10 min at room temperature, washed 2 times with ice-cold PBS, and stored at 4°C in PBS.

Image processing and analysis were performed by Fiji software^96^.

#### Chemically-inducible co-recruitment assay

EYFP-FKBP was fused to iSH2 or indicated mutants, while FRB-CFP is tethered to the inner leaflet of plasma membrane using the CAAX-region of K-Ras. Upon rapamycin addition, FKBP binds to FRB which brings the bait (mVenus-FKBP-iSH2) and the prey capable of binding (AP-2-mCherry or mCherry) to the plasma membrane. Recruitment of the bait and the prey to the plasma membrane were detected by TIRF microscopy as an increased fluorescence signal (Extended Data Fig. 6a–c). For quantification, after background subtraction, co-recruitment levels of prey were measured by increase in mCherry (prey) signal normalized to the intensity before rapamycin addition. Only cells showing at least 30% increase in mVenus (bait) intensity after Rapamycin addition were considered.

### Quantification and statistical analysis

All the quantified data were obtained from 3 or more independent experiments except for Extended Data Fig. 10d. To statistically compare a pair of data, wilcox.test was used in R as Wilcoxon rank sum test. To statistically compare multiple data, pSDCFlig (Asymptotic option) of NSM3 library was used in R as Steel-Dwass test.

### Quantification of iSH2 puncta index

Following the method described in Supplementary Figure 13 of a previous paper^97^, we created 5×5 median-filtered images of YF-iSH2 images and divided the raw image by the filtered images. iSH2 puncta index was measured by quantifying standard deviation of cytosolic region of the divided YF-iSH2 images. To avoid including intensity fluctuation caused by plasma membrane, regions of interest were manually drawn. We used Cos7 cells for the analysis of iSH2 mutants and variants since the cell showed more homogenous background (e.g., in the case of negative control YF) than HeLa cells.

### Western blot

3.6×10^5^ cells/well (6-well) were seeded ∼16 hours before experiment. The cells were serum-starved for 5–6 hours, stimulated as described in figure legends with 5% CO_2_ at 37°C. The reaction was stopped by directly replacing the culture media with 100 µL ice-cold RIPA buffer (Cell Signaling, 9806S) supplemented with cOmplete protease inhibitor (1×, Roche, 11873580001), 1 mM PMSF, and phosphatase inhibitors (1× for each, Sigma P5726 and P0044). Since cooling on ice was not sufficient to stop dephosphorylation, it was critical to immediately replace the media with RIPA buffer. Soluble fraction was collected as supernatant after centrifugation (14,000×g for 10 min at 4 °C) and the protein concentration was measured by Bradford assay. The samples were mixed with SDS-sample buffer, boiled at 95°C for 5 min, and separated on polyacrylamide gel. Proteins were transferred to methanol pre-treated PVDF membrane by using Criterion Blotter (BioRad, 1704070JA). The membrane was blocked by rocking in blocking buffer (3%BSA, 1×TBS) for 30–60 min at RT, stained with primary antibodies by rocking in antibody buffer (3%BSA, 1×TBS, 0.1% Tween 20, 0.1% NaN_3_) overnight at 4°C, washed (5 min×3 times) with TBS-T, stained with secondary antibodies in antibody buffer for 1 hours at rt, and washed again (5 min×3 times) with TBST. Fluorescent signals were detected by Typhoon or Pharos and analyzed by Fiji software^96^.

### Transferrin uptake assay

Transferrin uptake assay was performed by following the previous literature. Briefly, MEF cells were serum starved in the imaging media containing DMEM (Corning, 17-205-CV) and 1×Glutamax (Thermo Fisher Scientific, 35050061) for more than 2 hours and incubated with 250 µg/mL of Alexa Fluor 647- conjugated transferrin for indicated time. Cells were then chilled on ice, washed 3 times with PBS, washed 3 times with acid solution (0.2 M acetic acid, 0.5 M NaCl, pH 4.1), washed 3 times with PBS, fixed with 4% paraformaldehyde in PBS at room temperature for 10 minutes, and washed with 3 times with PBS. The amount of endocytosed transferrin was measured by quantifying cytosolic intensity of Alexa Fluor 647 in epi fluorescence images.

### Immunofluorescence

Immunofluorescence against vinculin was performed as follows. 25×10^3^ cells/well MEF cells were seeded on fibronectin-coated 8-well chambers and incubated overnight in DMEM supplemented with 10% FBS. Cells were then washed with PBS twice, fixed with 4% paraformaldehyde in PBS at room temperature for 15 minutes, washed again with PBS twice, permeabilized 0.1 % Triton X-100 in PBS at room temperature for 2.5 minutes, and blocked with blocking buffer (1% BSA in PBS) at room temperature for 30 minutes. Antibody against vinculin was used as ×500 dilution in the blocking buffer and the binding was performed at 4°C overnight. The secondary antibody Alexa Fluor 568-conjugated anti-Mouse IgG was used as ×1000 dilution in the blocking buffer and the binding was performed at room temperature for 1 hour. Each antibody binding steps were followed by 3 times of 5 minutes wash with TBST.

### Proliferation assay

For proliferation assay, 2.5–5×10^4^ cells were seeded on flasks, cultured in DMEM supplemented with 10% FBS for 50–72 hours, and the final number of cells were counted. Doubling time was calculated by Initial and final number of cells assuming the cell growth is exponential.

### Random migration assay

24-well plate were coated with 10 µg/mL fibronectin (5 µg/cm^2^) >30 min at 37°C. 1×10^4^ MEF cells were seeded and incubated in DMEM supplemented with 1% FBS for roughly 20 hours. Cells were washed once with fresh DMEM supplemented with 1% FBS and the media were replaced with DMEM supplemented with 10% FBS and 0.25 µg/mL Hoechst 33342. Cells were left in a 37°C and 5% CO_2_ incubator for 2 hours (Hoechst stain seemed to delay in the presence of fibronectin or collagen coating). Random migration was performed at 37°C and with 5% CO_2_ and humidity. Images were captured every 10 minutes for 16 hours through DAPI channel and phase contrast and analyzed by TrackMate^98^ plugin in Fiji software^96^.

### Chemotaxis

Chemotaxis assay was performed on µ-slide chemotaxis chambers (ibidi, 80326) by following manufacturer’s protocol. Briefly, 2.4×10^6^/mL WT MEF or 3.0×10^6^/mL p85 DKO and rescued MEF were seeded. After incubation at 37°C with 5% CO_2_ and 95% humidity for 2–3 hours, right reservoir was filled with imaging media supplemented with 1% FBS and 0.25 µg/mL Hoechst 33342 and left reservoir was filled with imaging media supplemented with 20% FBS and 0.25 µg/mL Hoechst 33342. The chamber was further incubated for 2 hours to allow the FBS gradient to be established. Chemotaxis was performed at 37°C with 5% CO_2_ and humidity. Images were captured every 10 minutes for 16 hours through DAPI channel and bright field and analyzed by TrackMate plugin^98^ in Fiji software^96^.

---

**Extended Data Figure 1:**
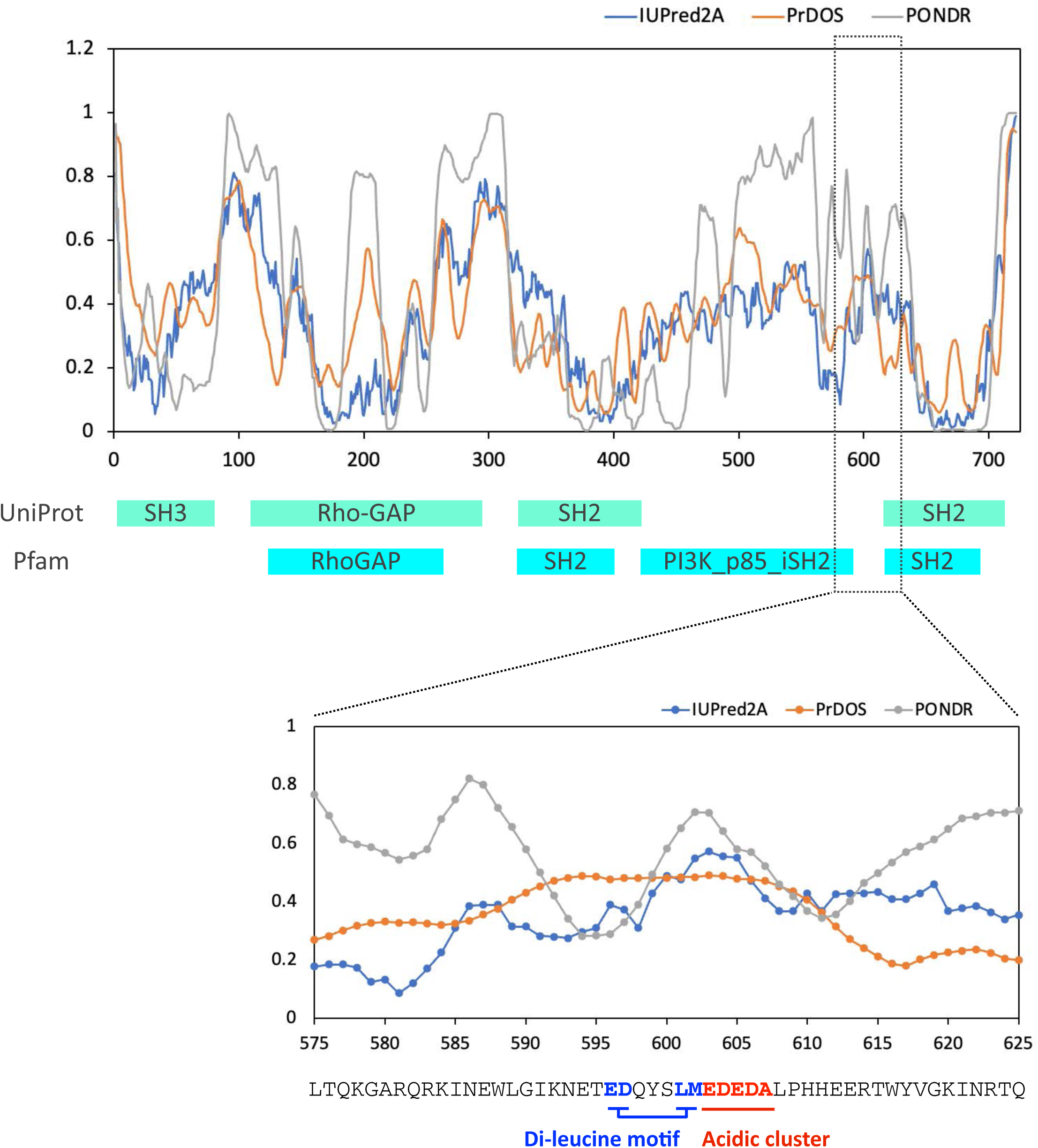
Prediction of intrinsically disordered regions. Intrinsically disordered region of mouse p85β (PIK3R2) was analyzed by three algorithms, IUPred2A, PrDOS, and PONDR.

**Extended Data Figure 2:**
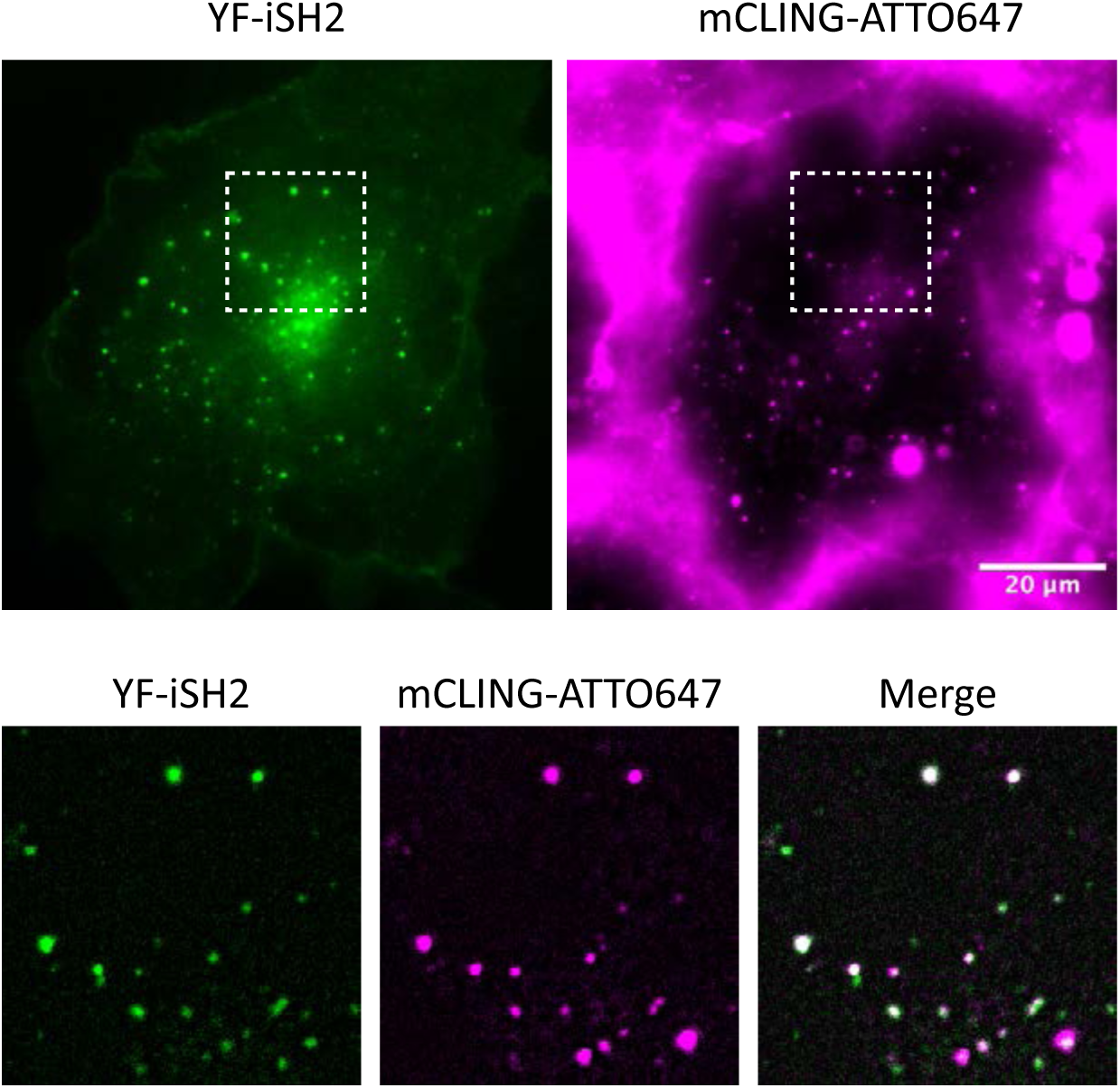
iSH2-vesicles colocalize with mCLING dye. Epi-fluorescence microscopy images of iSH2-vesicles colocalized with extracellularly added mCLING-ATTO647. Cos7 cells were transiently transfected with Lyn-ECFP-FRB, EYFP-FKBP-iSH2, mCherry-PH(Akt). After mCLING addition, iSH2 translocation and vesicle formation was induced by 100 nM rapamycin. 30 min after rapamycin addition, the samples were chilled, washed, and fixed with 4% paraformaldehyde. Top: raw image of a transfected cell. Bottom: enlarged images of dashed line area of top images. To reduce background noise, median filtered values were subtracted from the raw images.

**Extended Data Figure 3:**
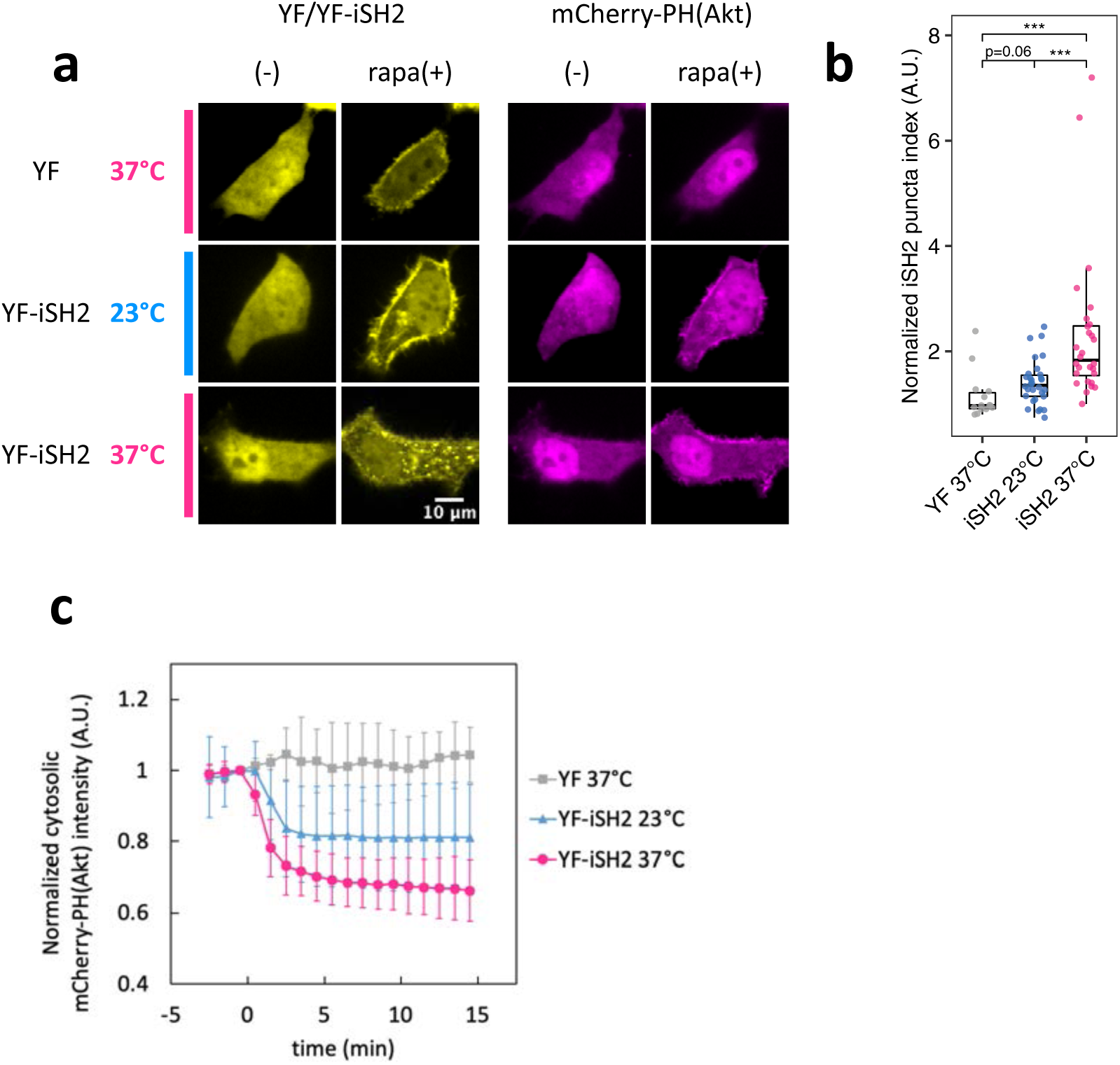
Temperature dependency of iSH2-mediated endocytosis. (a) Confocal images of endocytic vesicle production and PH(Akt) translocation. HeLa cells were transiently transfected with Lyn-ECFP-FRB, mCherry-PH(Akt), and EYFP-FKBP or EYFP-FKBP-iSH2. (-) before rapamycin addition, rapa(+) 20 min after adding 100 nM rapamycin. (b) Quantified iSH2-mediated endocytosis indices. The values were normalized by time=0. Box whisker plots represent median, 1st, 3rd quartiles and 1.5×inter- quartile range. P-value ***: < 0.001. Steel-Dwass test. (c) Time course of PH(Akt) translocation. Cytosolic intensity of mCherry-PH(Aki) was quantified and normalized by time=0. Error bars represent standard deviation. YF 37°C, n=15 cells. YF-iSH2 23°C, n=30 cells. YF-iSH2 37°C, n=28 cells.

**Extended Data Figure 4:**
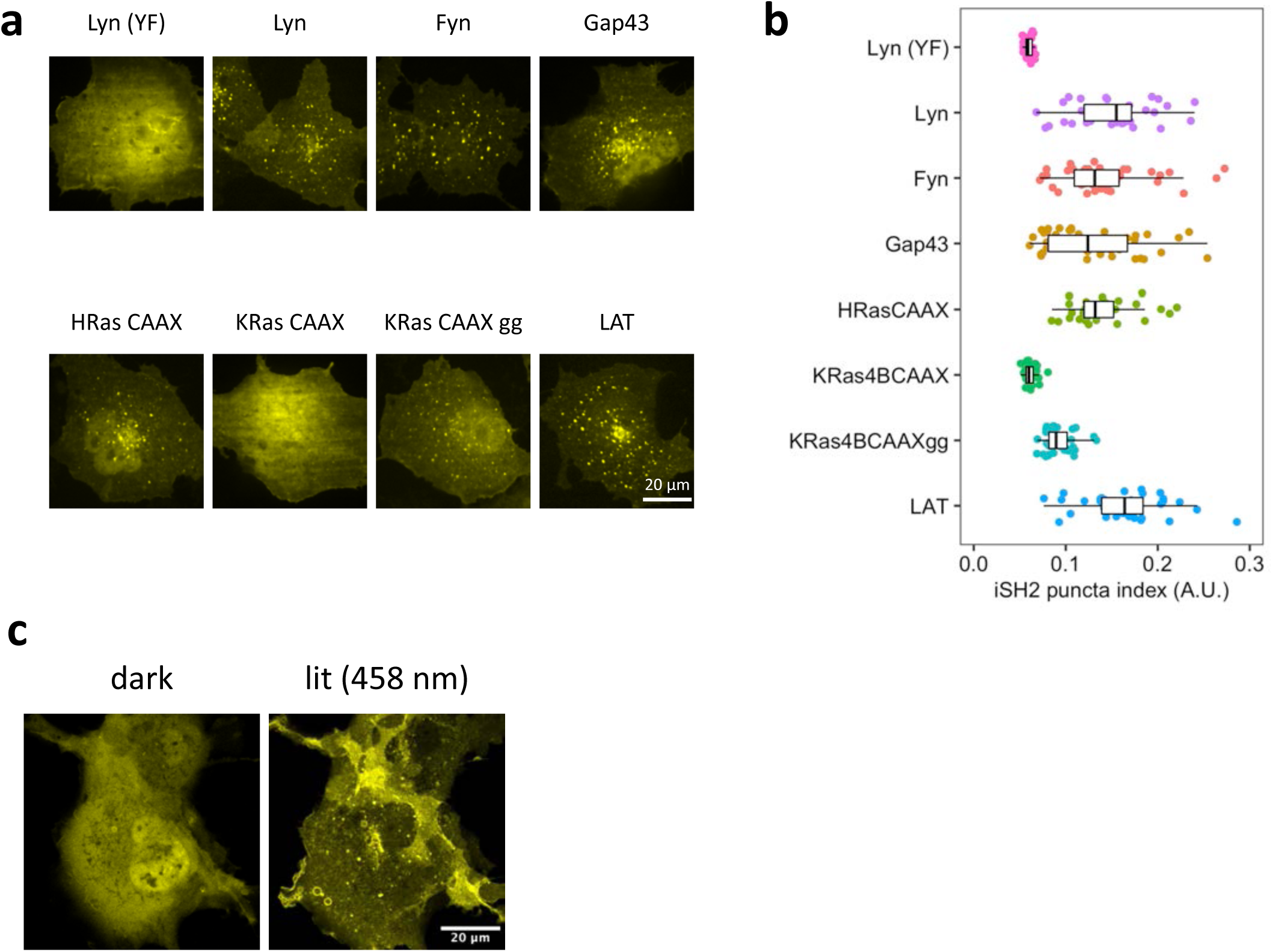
Generality of iSH2-mediated endocytosis. (a, b) Confocal images of iSH2- vesicles produced with different plasma membrane anchors and the quantified iSH2 puncta index. Cos7 cells were transiently transfected with EYFP-FKBP-iSH2, mCherry-PH(Akt), and ECFP-FRB fused with different types of plasma membrane anchors. 15 min after adding 100 nM rapamycin, cells were chilled, washed, and fixed with 4% paraformaldehyde and 0.15% glutaraldehyde. Box whisker plots represent median, 1st, 3rd quartiles and 1.5×inter-quartile range. (c) Confocal images of iSH2-vesicles induced by iLID/SspB system. Cos7 cells were transiently transfected with Lyn-iLID and EYFP-SspB-iSH2. dark: before light stimulation. lit (458 nm): 15 min after 458 nm light illumination. EYFP-SspB-iSH2 shows punctate structure in the cytosol.

**Extended Data Figure 5:**
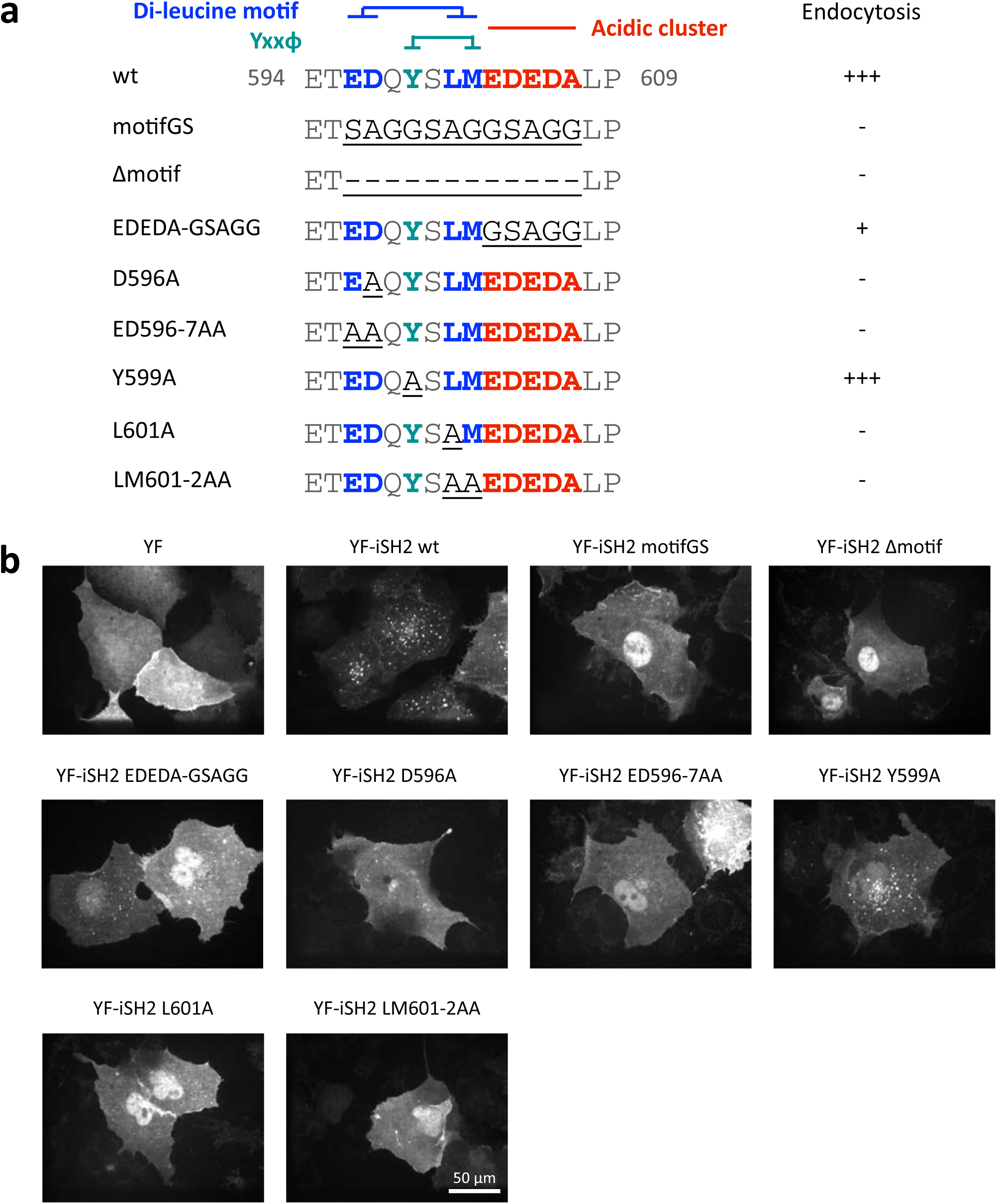
Vesicle formation with iSH2 variants. (a) List of the tested iSH2 mutants. Underlines indicate mutation sites. Here, wild type is derived from iSH2 domain of mouse p85β. (b) Confocal images of iSH2-vesicles produced with wild type and mutant iSH2. Cos7 cells were transiently transfected with Lyn-ECFP-FRB, EYFP-FKBP-iSH2, and mCherry-PH(Akt). 15 min after adding 100 nM rapamycin, cells were chilled, washed, and fixed with 4% paraformaldehyde and 0.15% glutaraldehyde.

**Extended Data Figure 6:**
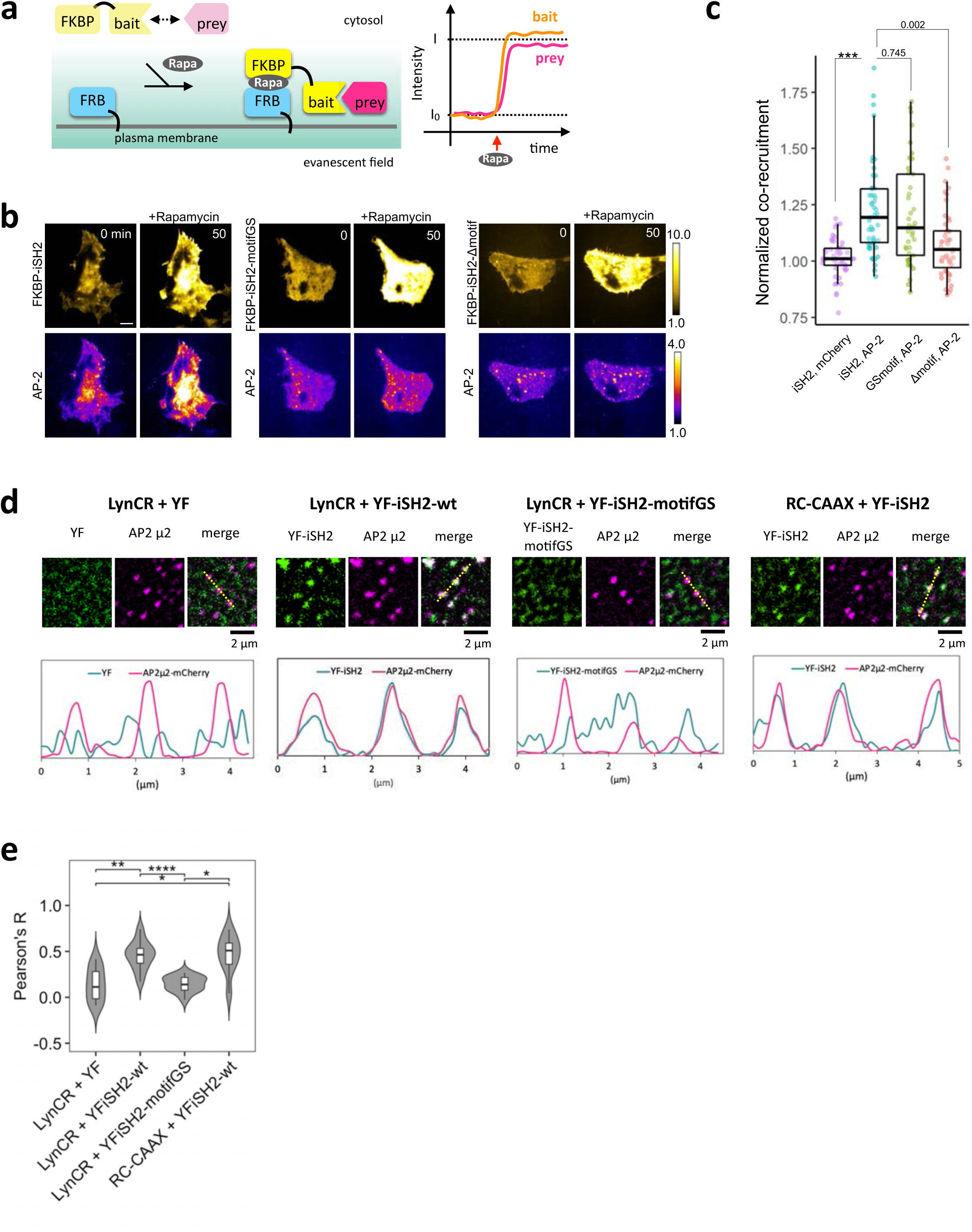
iSH2 recruits AP-2 to plasma membrane. (a) Schematic of co-recruitment assay. Interaction between bait and prey was evaluated by rapamycin-dependent increase in plasma membrane intensify of prey, here AP-2-mCherry. (b) Representative images showing changes in TIRF fluorescence intensities on plasma-membrane recruitment of EYFP-FKBP-iSH2, EYFP-FKBP-iSH2-motifGS and EYFP-FKBP-iSH2-Δmotif and corresponding changes in AP2-mCherry intensities. Scale bar: 10 µm. (c) Co-recruitment indices (I/I0) of mCherry with EYFP-FKBP-iSH2 and of AP2-mCherry with EYFP-FKBP-iSH2, EYFP-FKBP-iSH2-GSmotif and EYFP-FKBP-iSH2-Δmotif using the live cell co-recruitment assay. ***, P < 0.001 or as shown, Student’s t test. (d) TIRF images showing co-localization between YF-iSH2 and mCherry-AP-2(µ2). During live cell imaging, images were taken 1 min after 100 nM rapamycin addition. To reduce noise, median filtered images were subtracted from raw images. Graphs show line scan of dashed lines in merge images. (e) Pearson’s correlation between YFP signal and mCherry-AP-2(µ2) signal of (d). Calculation was performed on raw images. For each cell, 10 µm (80 pixels) × 10 µm (80 pixels) areas were selected for the quantification. Steel-Dwass test. P-values: *: < 0.05, **: < 0.01, ***: < 0.001, ****: < 0.0001.

**Extended Data Figure 7:**
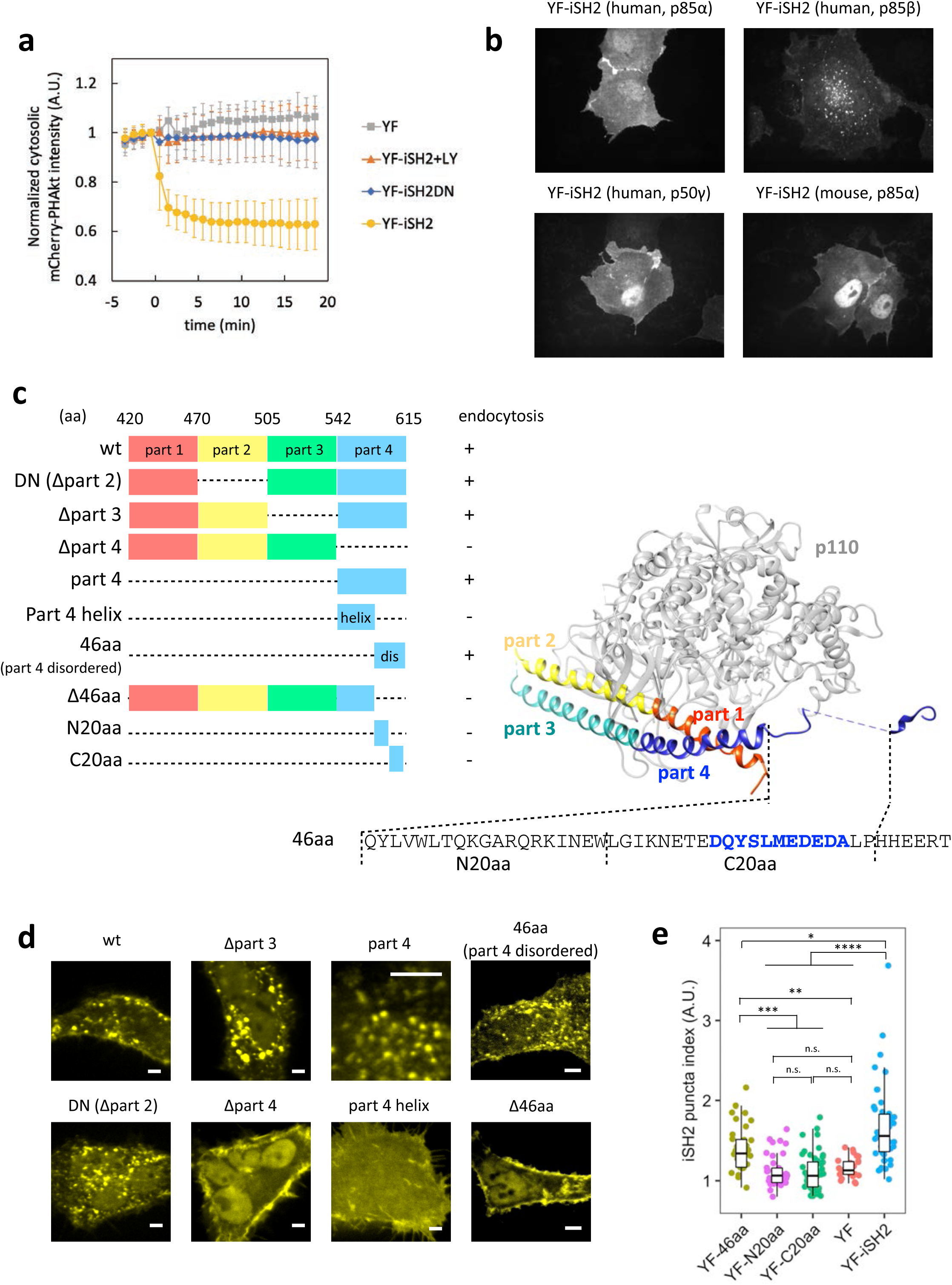
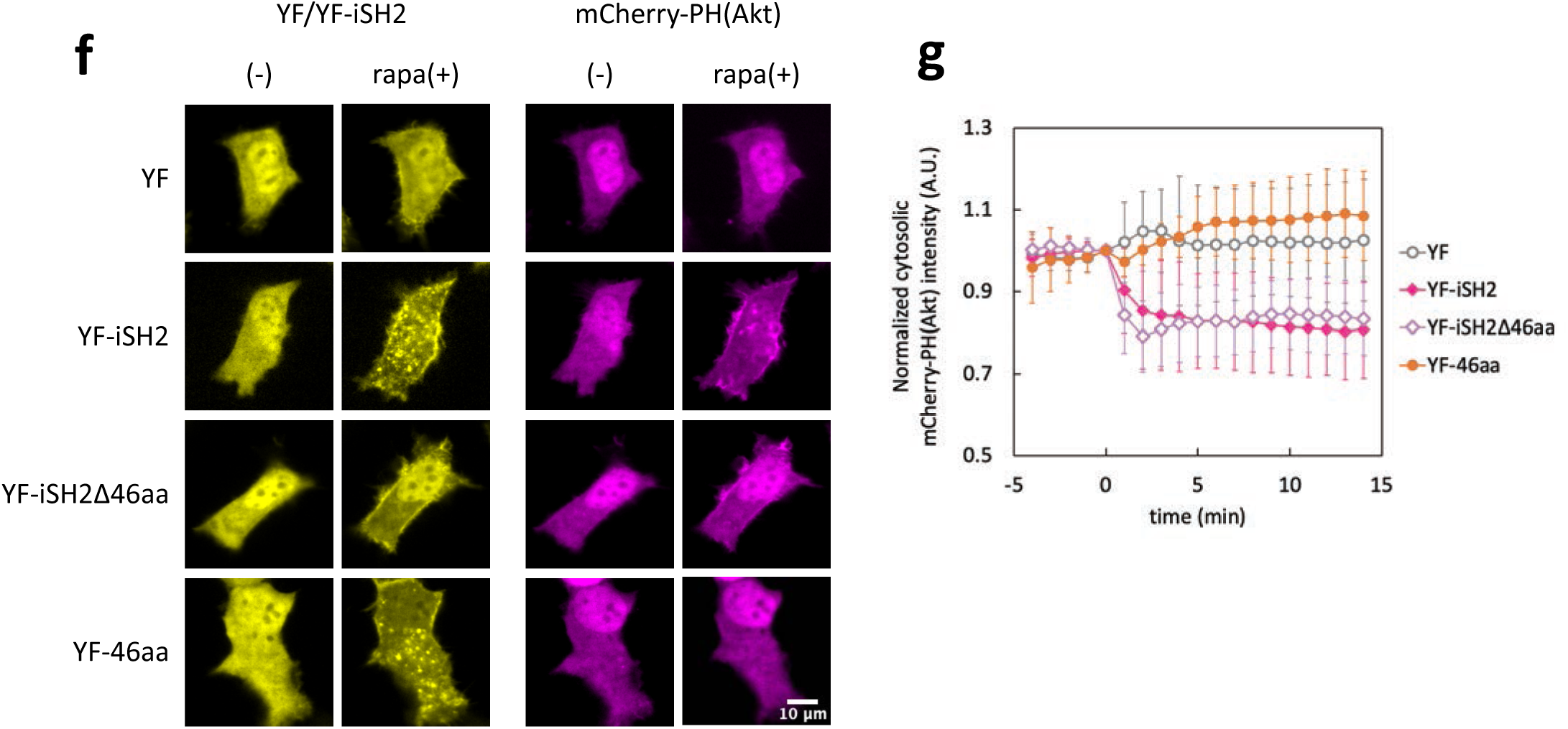
iSH2-mediated endocytosis is independent of PI3K catalytic activity and C- terminal 46 aa region is necessary and sufficient. (a) Time course of PH(Akt) translocation of Fig. 2a. Cytosolic intensity of mCherry-PH(Akt) was quantified and normalized by time=0. Error bars represent standard deviation. YF-iSH2, n=30 cells. YF-iSH2 + LY, n=28 cells. YF-iSH2DN, n=27 cells. YF, n=28 cells. (b) Confocal images of vesicles induced by iSH2 derived from different p85 isoforms. Cos7 cells were transiently transfected with Lyn-ECFP-FRB, EYFP-FKBP-iSH2, and mCherry-PH(Akt). 15 min after adding 100 nM rapamycin, cells were chilled, washed, and fixed with 4% paraformaldehyde and 0.15% glutaraldehyde. (c) Schematic representation of iSH2 truncates. Crystal structure of p110β-iSH2β is derived from PDB 2y3a. (d) Representative confocal image of live-cell plasma membrane recruitment of iSH2 truncates in HeLa expressing Lyn-ECFP-FRB, EYFP-FKBP-iSH2 (truncates), and mCherry-PH(Akt). Scale bar, 5 µm. (e) Quantified iSH2 puncta index of iSH2 truncates tested in Cos7 cells expressing Lyn- ECFP-FRB, EYFP-FKBP-iSH2 (truncates), and mCherry-PH(Akt). YF-46aa, n=38 cells. YF-N20aa, n=39 cells. YF-C20aa, n=48 cells. YF, n=27 cells. YF-iSH2, n=46 cells. Box whisker plots represent median, 1st, 3rd quartiles and 1.5×inter-quartile range. P-values (Steel-Dwass test): *: < 0.05, **: < 0.01, ***: < 0.001, ****: < 0.0001. n.s.: not significant. (f) Confocal live-cell images of iSH2-vesicles and PH(Akt) translocation. (g) Time course of PH(Akt) translocation of (f). Cytosolic intensity of mCherry-PH(Aki) was quantified and normalized by time=0. Error bars represent standard deviation. YF, n=17 cells. YF-iSH2, n=41 cells. YF-iSH2Δ46aa, n=39 cells. YF-46aa, n=22 cells. (f, g) Data correspond with Fig. 2c.

**Extended Data Figure 8:**
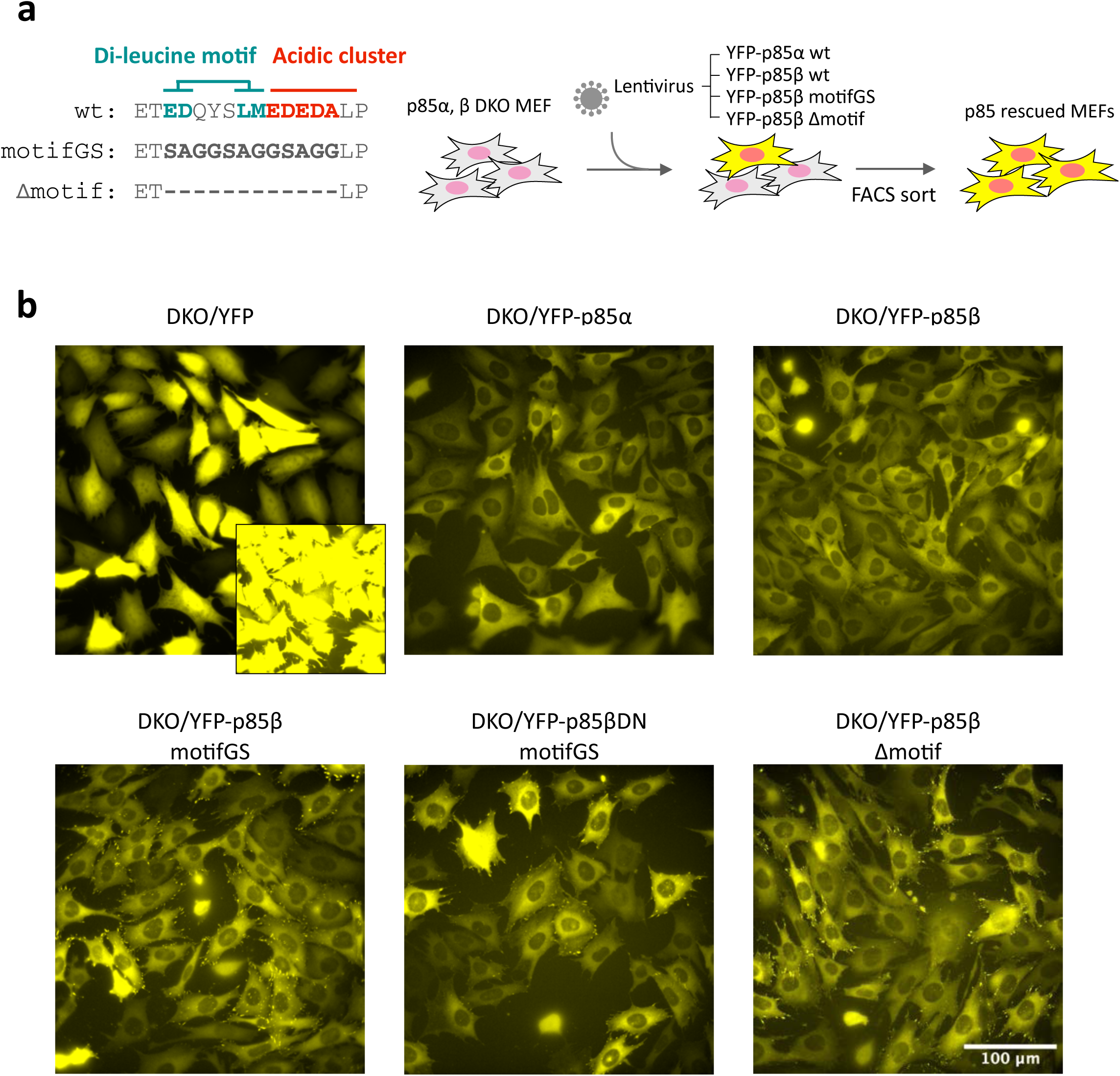
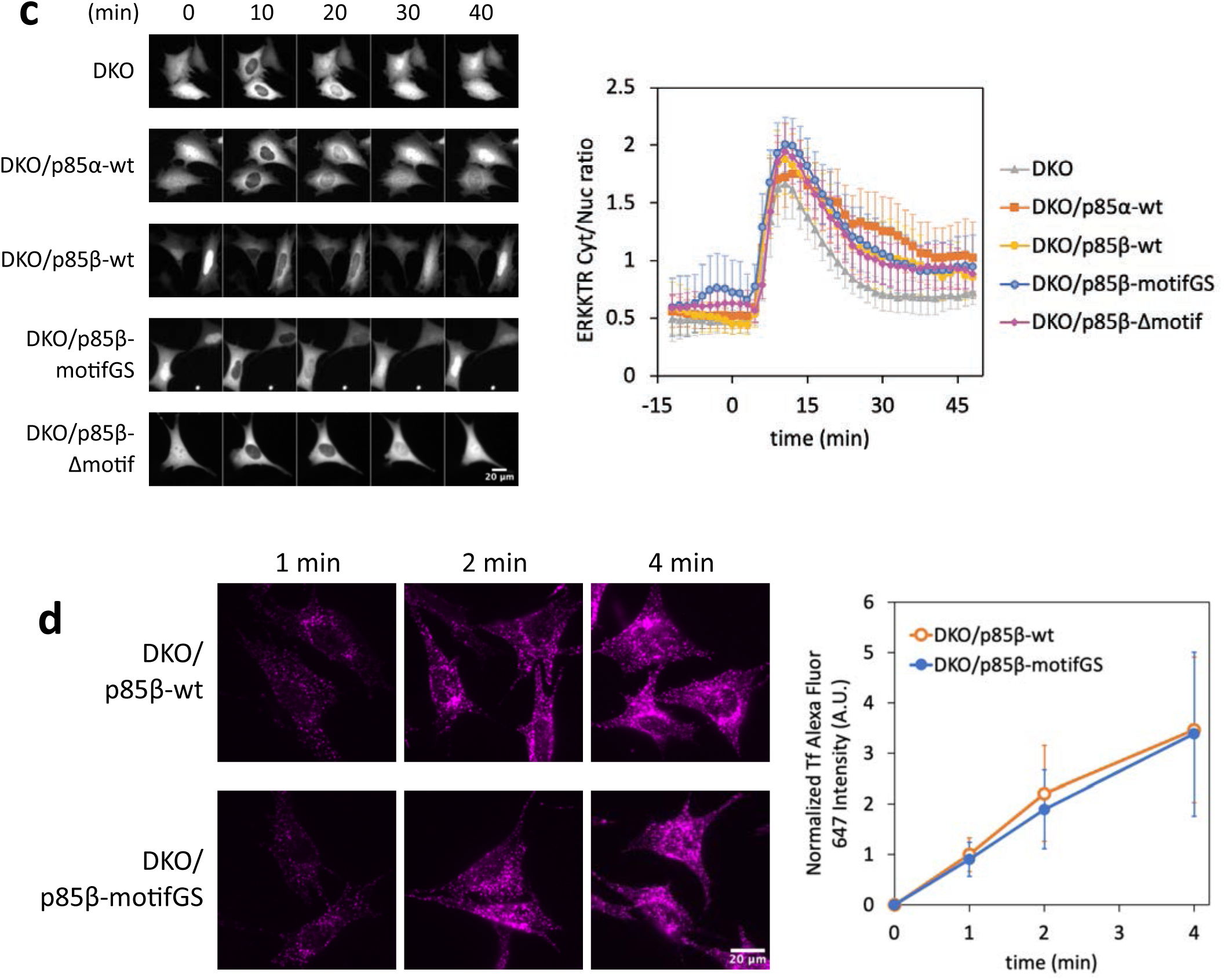
Generation and Functional analysis of p85-rescued MEFs. (a) p85α, β double knockout (DKO) MEFs were infected with lentiviruses encoding YFP-p85 variants. Infected cells were FACS-sorted by YFP fluorescence. (b) Epi-fluorescence microscopy images of each cell lines. Dynamic range was adjusted between. c) ERK response to PDGF stimulation. Each cell lines were transiently transfected with mCherry-ERKKTR. The cells were serum starved and stimulated with 50 ng/mL PDGF- BB. ERKKTR response was recorded by live cell imaging at 37°C with 5% CO_2_. Left: epi-fluorescence microscopy images of mCherry-ERKKTR. Right: quantified Cytosol/Nucleus ratio of mCherry-ERKKTR. Error bars represent 2×SEM (95% CI). DKO, n=18 cells. DKO/p85α-wt, n=18 cells. DKO/p85β-wt, n=18 cells. DKO/p85β-motifGS, n=19 cells. DKO/p85β-Δmotif, n=19 cells. (d) Transferrin uptake. Alexa Fluor 647-conjugated transferrin was added to serum starved cells. After the indicated time, the cells were chilled, washed with acid, and fixed with 4% paraformaldehyde. Left: epi-fluorescence microscopy images of Alexa Fluor 647-conjugated transferrin. Right: quantified Alexa Fluor 647 intensity. Error bars represent standard deviation. n>61 cells for each time point.

**Extended Data Figure 9:**
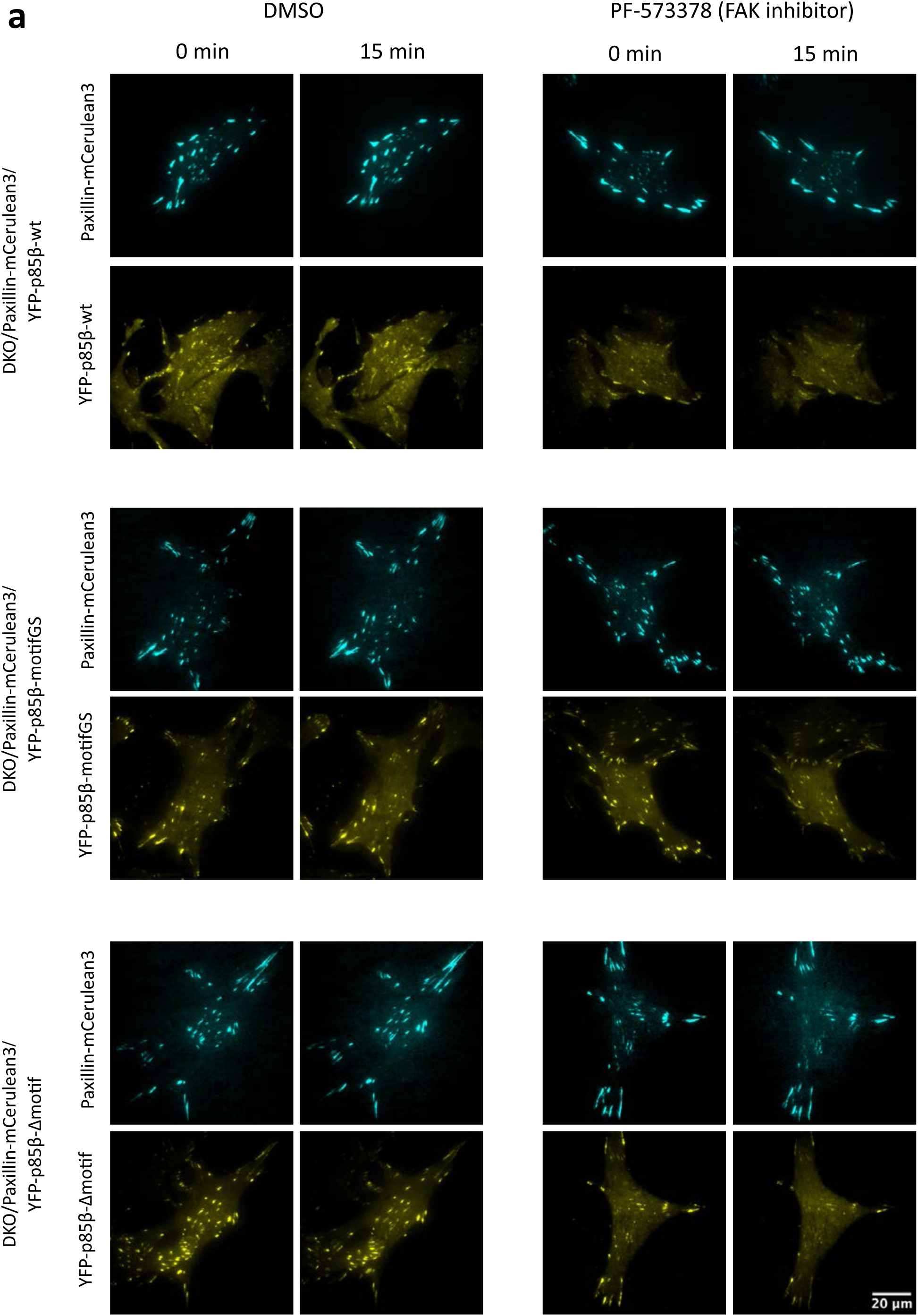
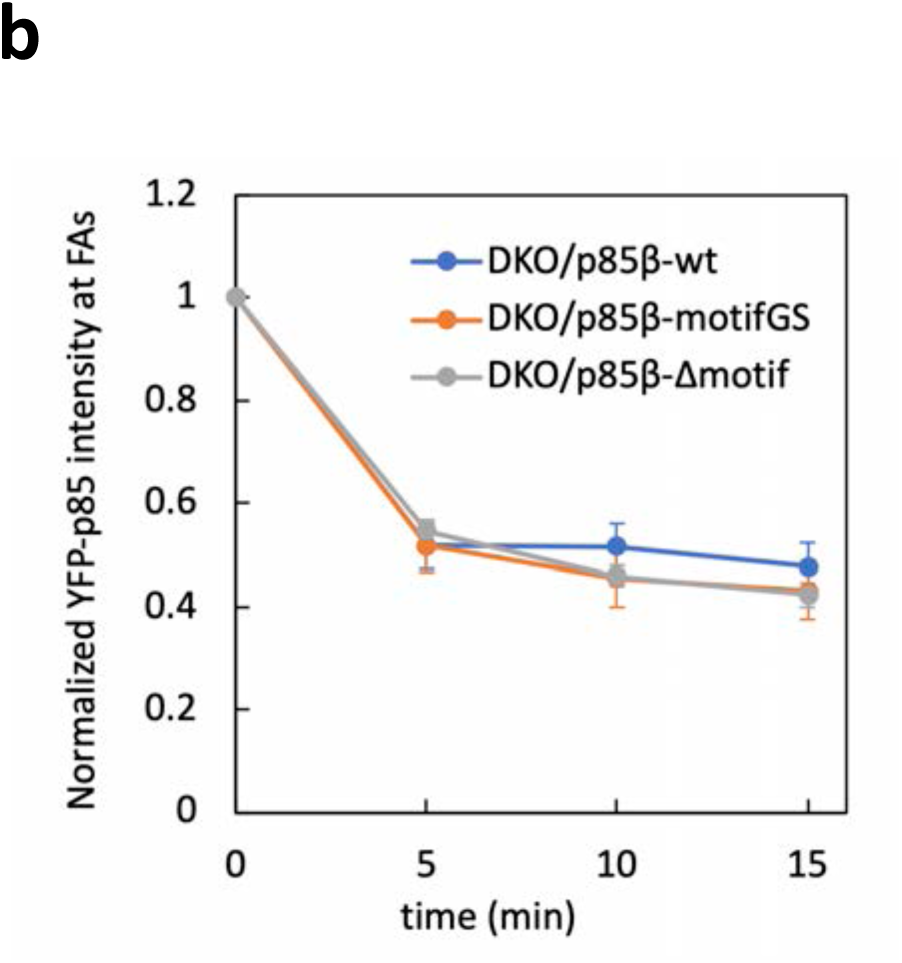
PF-573378 (FAK inhibitor) response of p85 variants. (a) TIRF images of MEFs stably expressing Paxillin-mCerulean3 and YFP-p85 variants. The cells were serum starved and imaged at 37°C with 5% CO_2_. (b) Normalized YFP-p85 intensity at focal adhesions. YFP-p85 intensity at focal adhesion was measured with image masks created by Paxillin-mCerulean3 images and normalized by time=0. Error bars represent standard deviation. DKO/p85β-wt, n=20 cells. DKO/p85β-motifGS, n=22 cells. DKO/p85β-Δmotif, n=18 cells.

**Extended Data Figure 10:**
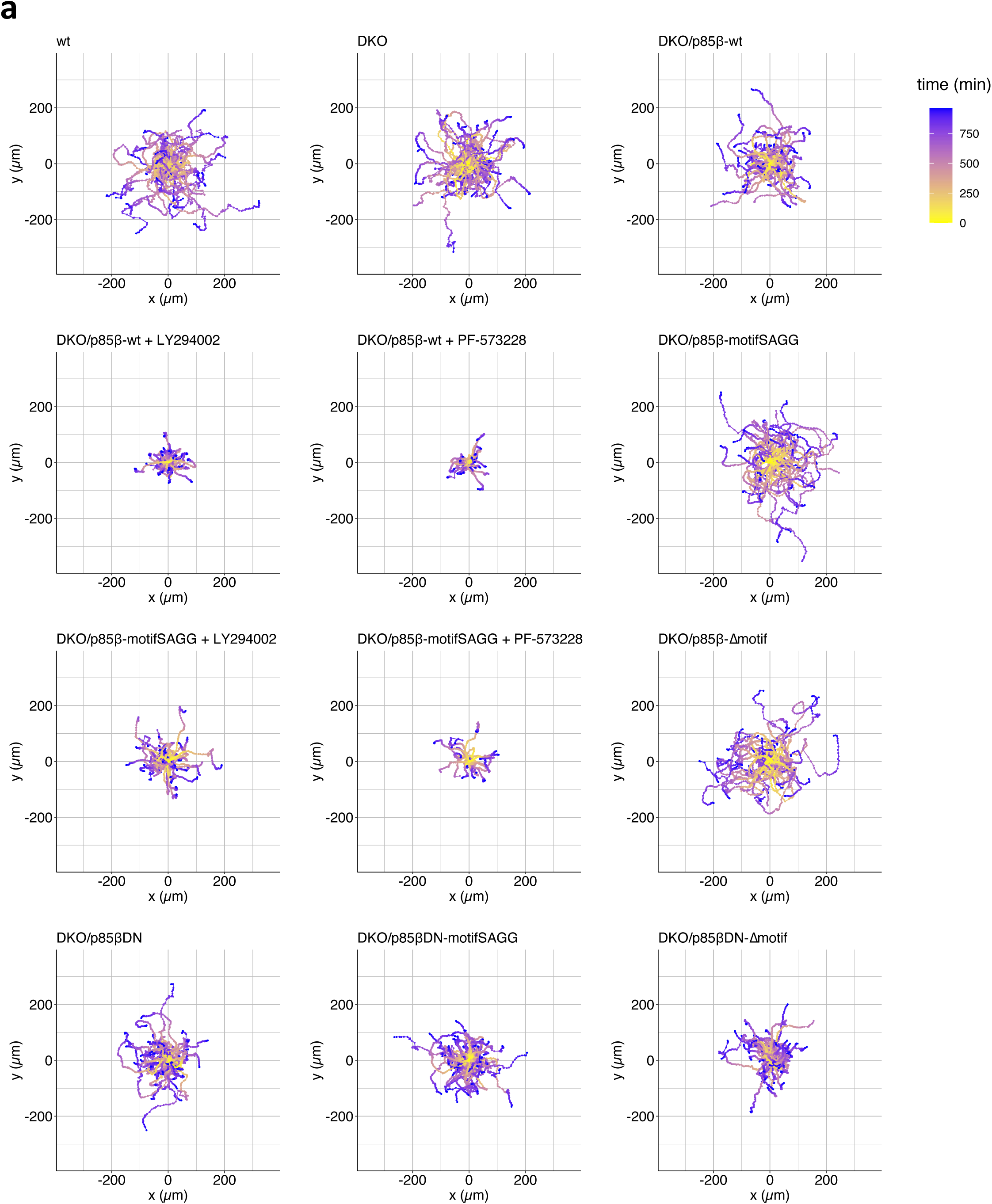
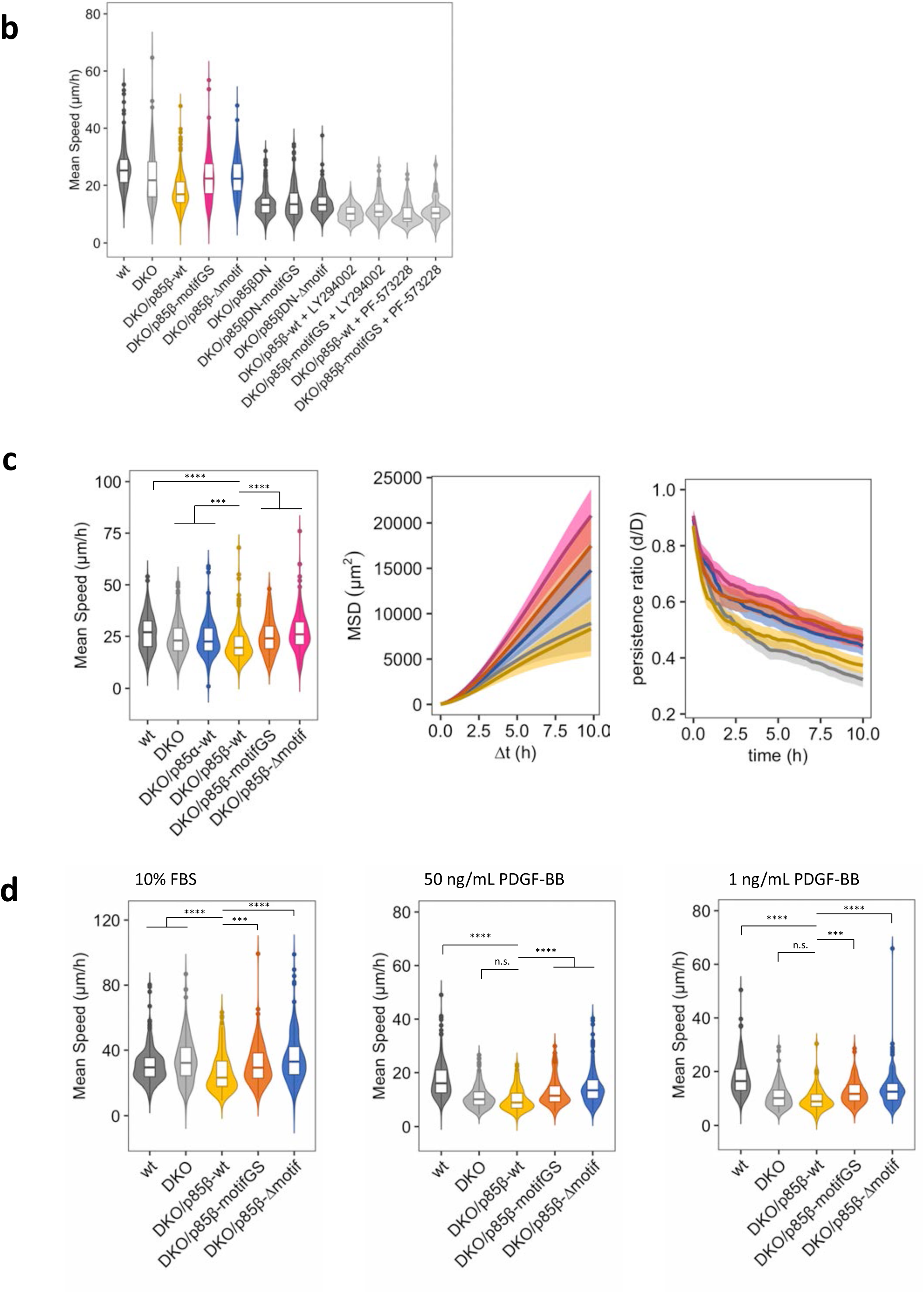
Supplementary data of migration assay. (a–c) Random migration. (a) Cell track analysis of each cell lines. Data correspond with Fig. 4b-d. (b) Full data of random migration including PI3K inhibitor LY294002 data and FAK inhibitor PF-573228 data. Data correspond with Fig. 4b-d. (c) Different data set of random migration including DKO/p85α-wt.

**Supplementary movie 1:** Confocal images of endocytic vesicles produced by plasma membrane targeting of iSH2 domain. HeLa cells were transiently transfected with Lyn-ECFP-FRB, EYFP-FKBP-iSH2, and mCherry-PH(Akt). Imaging was performed at 37°C with 5%CO_2_. 100 nM rapamycin was added at indicated time.

**Supplementary movie 2:** Confocal images of EYFP-FKBP negative control. HeLa cells were transiently transfected with Lyn-ECFP-FRB, EYFP-FKBP, and mCherry-PH(Akt). Imaging was performed at 37°C with 5%CO_2_. 100 nM rapamycin was added at indicated time.

**Supplementary movie 3**: Confocal images of room temperature control. HeLa cells were transiently transfected with Lyn-ECFP-FRB, EYFP-FKBP-iSH2, and mCherry-PH(Akt). Imaging was performed at 23°C with 5%CO_2_. 100 nM rapamycin was added at indicated time.

## References

1. Vanhaesebroeck, B., Guillermet-Guibert, J., Graupera, M. & Bilanges, B. The emerging mechanisms of isoform-specific PI3K signalling. Nature Reviews Molecular Cell Biology 11, 329–341 (2010).

2. Fruman, D. A. et al. The PI3K Pathway in Human Disease. Cell 170, 605–635 (2017).

3. Marat, A. L. & Haucke, V. Phosphatidylinositol 3-phosphates—at the interface between cell signalling and membrane traffic. The EMBO Journal 35, 561–579 (2016).

4. Bilanges, B., Posor, Y. & Vanhaesebroeck, B. PI3K isoforms in cell signalling and vesicle trafficking. Nature Reviews Molecular Cell Biology 20, 515–534 (2019).

5. Bear, J. E. & Haugh, J. M. Directed migration of mesenchymal cells: Where signaling and the cytoskeleton meet. Current Opinion in Cell Biology 30, 74–82 (2014).

6. Fang, D. & Liu, Y. C. Proteolysis-independent regulation of P13K by Cbl-b-mediated ubiquitination in T cells. Nature Immunology 2, 870–875 (2001).

7. Chiu, Y.-H., Lee, J. Y. & Cantley, L. C. BRD7, a Tumor Suppressor, Interacts with p85α and Regulates PI3K Activity. Molecular Cell 54, 193–202 (2014).

8. Furuya, F., Ying, H., Zhao, L. & Cheng, S. Novel functions of thyroid hormone receptor mutants: Beyond nucleus-initiated transcription. Steroids 72, 171–179 (2007).

9. Tsuboi, N. et al. The tyrosine phosphatase CD148 interacts with the p85 regulatory subunit of phosphoinositide 3-kinase. Biochemical Journal 413, 193–200 (2008).

10. Thapa, N. et al. Phosphatidylinositol 3-kinase Signaling is Spatially Organized at Endosomal 1 Compartments by Microtubule-associated Protein 4. Nature Cell Biology 22, 1357–1370 (2020).

11. Fruman, D. A. Regulatory Subunits of Class IA PI3K. in Phosphoinositide 3-kinase in Health and Disease (eds. Rommel, C., Vanhaesebroeck, B. & Vogt, P. K.) vol. 346 225–244 (Springer Berlin Heidelberg, 2010).

12. Tsolakos, N. et al. Quantitation of class IA PI3Ks in mice reveals p110-free-p85s and isoform- selective subunit associations and recruitment to receptors. Proceedings of the National Academy of Sciences of the United States of America 115, 12176–12181 (2018).

13. Fantl, W. J. et al. Distinct phosphotyrosines on a growth factor receptor bind to specific molecules that mediate different signaling pathways. Cell 69, 413–423 (1992).

14. Valius, M. & Kazlauskas, A. Phospholipase C-γ1 and phosphatidylinositol 3 kinase are the downstream mediators of the PDGF receptor’s mitogenic signal. Cell 73, 321–334 (1993).

15. Yu, J. et al. Regulation of the p85/p110 Phosphatidylinositol 3ʹ-Kinase: Stabilization and Inhibition of the p110α Catalytic Subunit by the p85 Regulatory Subunit. Molecular and Cellular Biology 18, 1379–1387 (1998).

16. Zhang, X. et al. Structure of Lipid Kinase p110β/p85β Elucidates an Unusual SH2-Domain- Mediated Inhibitory Mechanism. Molecular Cell 41, 567–578 (2011).

17. Parent, C. A., Blacklock, B. J., Froehlich, W. M., Murphy, D. B. & Devreotes, P. N. G Protein Signaling Events Are Activated at the Leading Edge of Chemotactic Cells. Cell 95, 81–91 (1998).

18. Meili, R. Chemoattractant-mediated transient activation and membrane localization of Akt/PKB is required for efficient chemotaxis to cAMP in Dictyostelium. The EMBO Journal 18, 2092–2105 (1999).

19. Servant, G. et al. Polarization of chemoattractant receptor signaling during neutrophil chemotaxis. Science 287, 1037–1040 (2000).

20. Weiner, O. D. et al. A PtdInsP3- and Rho GTPase-mediated positive feedback loop regulates neutrophil polarity. Nat Cell Biol 4, 509–513 (2002).

21. Srinivasan, S. et al. Rac and Cdc42 play distinct roles in regulating PI(3,4,5)P3 and polarity during neutrophil chemotaxis. Journal of Cell Biology 160, 375–385 (2003).

22. Schneider, I. C. & Haugh, J. M. Quantitative elucidation of a distinct spatial gradient-sensing mechanism in fibroblasts. Journal of Cell Biology 171, 883–892 (2005).

23. Welf, E. S., Ahmed, S., Johnson, H. E., Melvin, A. T. & Haugh, J. M. Migrating fibroblasts reorient directionality by a metastable, PI3K-dependent mechanism. Journal of Cell Biology 197, 105–114 (2012).

24. Johnson, H. E. et al. F-actin bundles direct the initiation and orientation of lamellipodia through adhesion-based signaling. Journal of Cell Biology 208, 443–455 (2015).

25. Park, S. W. et al. The regulatory subunits of PI3K, p85α and p85β, interact with XBP-1 and increase its nuclear translocation. Nat Med 16, 429–437 (2010).

26. Winnay, J. N., Boucher, J., Mori, M. A., Ueki, K. & Kahn, C. R. A regulatory subunit of phosphoinositide 3-kinase increases the nuclear accumulation of X-box–binding protein-1 to modulate the unfolded protein response. Nat Med 16, 438–445 (2010).

27. Luo, J., Field, S. J., Lee, J. Y., Engelman, J. A. & Cantley, L. C. The p85 regulatory subunit of phosphoinositide 3-kinase down-regulates IRS-1 signaling via the formation of a sequestration complex. Journal of Cell Biology 170, 455–464 (2005).

28. Chamberlain, M. D. et al. Deregulation of Rab5 and Rab4 proteins in p85R274A-expressing cells alters PDGFR trafficking. Cellular Signalling 22, 1562–1575 (2010).

29. Bulut, G. B., Sulahian, R., Yao, H. & Huang, L. J. S. Cbl ubiquitination of p85 is essential for Epo-induced EpoR endocytosis. Blood 122, 3964–3972 (2013).

30. Jiménez, C. et al. Role of the Pi3k Regulatory Subunit in the Control of Actin Organization and Cell Migration. Journal of Cell Biology 151, 249–262 (2000).

31. García, Z. et al. A PI3K activity-independent function of p85 regulatory subunit in control of mammalian cytokinesis. EMBO J 25, 4740–4751 (2006).

32. Kumar, M. et al. The Eukaryotic Linear Motif resource: 2022 release. Nucleic Acids Research 50, D497–D508 (2022).

33. Traub, L. M. & Bonifacino, J. S. Cargo recognition in clathrin-mediated endocytosis. Cold Spring Harbor Perspectives in Biology 5, 1–24 (2013).

34. Mészáros, B., Erdős, G. & Dosztányi, Z. IUPred2A: context-dependent prediction of protein disorder as a function of redox state and protein binding. Nucleic Acids Research 46, W329–W337 (2018).

35. Ishida, T. & Kinoshita, K. PrDOS: prediction of disordered protein regions from amino acid sequence. Nucleic Acids Research 35, W460–W464 (2007).

36. Predictor of Natural Disordered Regions (PONDR). http://www.pondr.com/.

37. Terrillon, S. & Bouvier, M. Receptor activity-independent recruitment of βarrestin2 reveals specific signalling modes. EMBO J 23, 3950–3961 (2004).

38. Wood, L. A., Larocque, G., Clarke, N. I., Sarkar, S. & Royle, S. J. New tools for ‘hot-wiring’ clathrin- mediated endocytosis with temporal and spatial precision. Journal of Cell Biology 216, 4351–4365 (2017).

39. DeRose, R., Miyamoto, T. & Inoue, T. Manipulating signaling at will: chemically-inducible dimerization (CID) techniques resolve problems in cell biology. Pflugers Archiv : European journal of physiology 465, 409–417 (2013).

40. Revelo, N. H. et al. A new probe for super-resolution imaging of membranes elucidates trafficking pathways. Journal of Cell Biology 205, 591–606 (2014).

41. Leonard, M., Doo Song, B., Ramachandran, R. & Schmid, S. L. Robust Colorimetric Assays for Dynamin’s Basal and Stimulated GTPase Activities. in Methods in Enzymology vol. 404 490–503 (Elsevier, 2005).

42. Delvendahl, I., Vyleta, N. P., von Gersdorff, H. & Hallermann, S. Fast, Temperature-Sensitive and Clathrin-Independent Endocytosis at Central Synapses. Neuron 90, 492–498 (2016).

43. Suh, B. C., Inoue, T., Meyer, T. & Hille, B. Rapid chemically induced changes of PtdIns(4,5)P2 gate KCNQ ion channels. Science 314, 1454–1457 (2006).

44. Inoue, T. & Meyer, T. Synthetic Activation of Endogenous PI3K and Rac Identifies an AND-Gate Switch for Cell Polarization and Migration. PLOS ONE 3, e3068 (2008).

45. Nakatsu, F. et al. The inositol 5-phosphatase SHIP2 regulates endocytic clathrin-coated pit dynamics. Journal of Cell Biology 190, 307–315 (2010).

46. Thevathasan, J. V. et al. The small GTPase HRas shapes local PI3K signals through positive feedback and regulates persistent membrane extension in migrating fibroblasts. MBoC 24, 2228–2237 (2013).

47. Guntas, G. et al. Engineering an improved light-induced dimer (iLID) for controlling the localization and activity of signaling proteins. Proceedings of the National Academy of Sciences of the United States of America 112, 112–117 (2015).

48. Cilleros-Rodriguez, D. et al. Multiple ciliary localization signals control INPP5E ciliary targeting. eLife 11, e78383 (2022).

49. Deb Roy, A., et al. Non-catalytic allostery in α-TAT1 by a phospho-switch drives dynamic microtubule acetylation. Journal of Cell Biology 221, e202202100 (2022).

50. Ford, M. G. J. et al. Simultaneous binding of PtdIns (4,5) P2 and clathrin by AP180 in the nucleation of clathrin lattices on membranes. Science 291, 1051–1055 (2001).

51. Zhao, X. et al. Expression of auxilin or AP180 inhibits endocytosis by mislocalizing clathrin: Evidence for formation of nascent pits containing AP1 or AP2 but not clathrin. Journal of Cell Science 114, 353–365 (2001).

52. van der Bliek, A. et al. Mutations in human dynamin block an intermediate stage in coated vesicle formation. Journal of Cell Biology 122, 553–563 (1993).

53. Damke, H., Baba, T., Warnock, D. E. & Schmid, S. L. Induction of mutant dynamin specifically blocks endocytic coated vesicle formation. Journal of Cell Biology 127, 915–934 (1994).

54. Basquin, C. et al. The signalling factor PI3K is a specific regulator of the clathrin-independent dynamin-dependent endocytosis of IL-2 receptors. Journal of Cell Science 126, 1099–1108 (2013).

55. Laketa, V. et al. PIP3 induces the recycling of receptor tyrosine kinases. Science Signaling 7, 1– 10 (2014).

56. Boucrot, E. et al. Endophilin marks and controls a clathrin-independent endocytic pathway. Nature 517, 460–465 (2015).

57. Goulden, B. D. et al. A high-avidity biosensor reveals plasma membrane PI(3,4)P2 is predominantly a class I PI3K signaling product. Journal of Cell Biology 218, 1066–1079 (2019).

58. Walker, E. H. et al. Structural Determinants of Phosphoinositide 3-Kinase Inhibition by Wortmannin, LY294002, Quercetin, Myricetin, and Staurosporine. Molecular Cell 11, 909–919 (2000).

59. Dhand, R. et al. PI 3-kinase: Structural and functional analysis of intersubunit interactions. EMBO Journal 13, 511–521 (1994).

60. Hale, B. G., Jackson, D., Chen, Y.-H., Lamb, R. A. & Randall, R. E. Influenza A virus NS1 protein binds p85β and activates phosphatidylinositol-3-kinase signaling. Proc. Natl. Acad. Sci. U.S.A. 103, 14194–14199 (2006).

61. Li, Y., Anderson, D. H., Liu, Q. & Zhou, Y. Mechanism of Influenza A Virus NS1 Protein Interaction with the p85β, but Not the p85α, Subunit of Phosphatidylinositol 3-Kinase (PI3K) and Up-regulation of PI3K Activity. Journal of Biological Chemistry 283, 23397–23409 (2008).

62. Hale, B. G. et al. Structural insights into phosphoinositide 3-kinase activation by the influenza A virus NS1 protein. Proceedings of the National Academy of Sciences of the United States of America 107, 1954–1959 (2010).

63. Brachmann, S. M. et al. Role of Phosphoinositide 3-Kinase Regulatory Isoforms in Development and Actin Rearrangement. Molecular and Cellular Biology 25, 2593–2606 (2005).

64. Brüggemann, Y., Karajannis, L. S., Stanoev, A., Stallaert, W. & Bastiaens, P. I. H. Growth factor– dependent ErbB vesicular dynamics couple receptor signaling to spatially and functionally distinct Erk pools. Sci. Signal. 14, eabd9943 (2021).

65. Cariaga-Martinez, A. E. et al. Phosphoinositide 3-kinase p85beta regulates invadopodium formation. Biology Open 3, 924–936 (2014).

66. Reiske, H. R. et al. Requirement of Phosphatidylinositol 3-Kinase in Focal Adhesion Kinase- promoted Cell Migration. Journal of Biological Chemistry 274, 12361–12366 (1999).

67. Chen, H. C. & Guan, J. L. Association of focal adhesion kinase with its potential substrate phosphatidylinositol 3-kinase. Proc. Natl. Acad. Sci. U.S.A. 91, 10148–10152 (1994).

68. Chen, H. C., Appeddu, P. A., Isoda, H. & Guan, J. L. Phosphorylation of tyrosine 397 in focal adhesion kinase is required for binding phosphatidylinositol 3-kinase. Journal of Biological Chemistry 271, 26329–26334 (1996).

69. Bachelot, C., Rameh, L., Parsons, T. & Cantley, L.C. Association of phosphatidylinositol 3-kinase, via the SH2 domains of p85, with focal adhesion kinase in polyoma middle t-transformed fibroblasts. Biochim Biophys Acta 1311, 45–52 (1996).

70. Gillham, H., Golding, M. C. H. M., Pepperkok, R. & Gullick, W. J. Intracellular movement of green fluorescent protein-tagged phosphatidylinositol 3-kinase in response to growth factor receptor signaling. Journal of Cell Biology 146, 869–880 (1999).

71. Stehbens, S. J. & Wittmann, T. Analysis of focal adhesion turnover. A quantitative live-cell imaging example. Methods in Cell Biology vol. 123 (Elsevier Inc., 2014).

72. Slack-Davis, J. K. et al. Cellular Characterization of a Novel Focal Adhesion Kinase Inhibitor. Journal of Biological Chemistry 282, 14845–14852 (2007).

73. Case, L. B. & Waterman, C. M. Integration of actin dynamics and cell adhesion by a three- dimensional, mechanosensitive molecular clutch. Nat Cell Biol 17, 955–963 (2015).

74. Hu, Q., Klippel, A., Muslin, A. J., Fantl, W. J. & Williams, L. T. Ras-dependent induction of cellular responses by constitutively active phosphatidylinositol-3 kinase. Science 268, 100–102 (1995).

75. Sun, J., Singaram, I., Soflaee, M. H. & Cho, W. A direct fluorometric activity assay for lipid kinases and phosphatases. Journal of Lipid Research 61, 945–952 (2020).

76. Taylor, M. J., Perrais, D. & Merrifield, C. J. A high precision survey of the molecular dynamics of mammalian clathrin-mediated endocytosis. PLoS Biology 9, (2011).

77. Devreotes, P. N. et al. Excitable Signal Transduction Networks in Directed Cell Migration. Annu. Rev. Cell Dev. Biol. 33, 103–125 (2017).

78. Weiger, M. C., Ahmed, S., Welf, E. S. & Haugh, J. M. Directional persistence of cell migration coincides with stability of asymmetric intracellular signaling. Biophysical Journal 98, 67–75 (2010).

79. Vallejo-Díaz, J., Chagoyen, M., Olazabal-Morán, M., González-García, A. & Carrera, A. C. The Opposing Roles of PIK3R1/p85α and PIK3R2/p85β in Cancer. Trends in Cancer 5, 233–244 (2019).

80. Cortés, I. et al. P85Β Phosphoinositide 3-Kinase Subunit Regulates Tumor Progression. Proceedings of the National Academy of Sciences of the United States of America 109, 11318–11323 (2012).

81. Ito, Y., Hart, J. R., Ueno, L. & Vogt, P. K. Oncogenic activity of the regulatory subunit p85β of phosphatidylinositol 3-kinase (PI3K). PNAS 111, 16826–16829 (2014).

82. Ito, Y., Vogt, P. K. & Hart, J. R. Domain analysis reveals striking functional differences between the regulatory subunits of phosphatidylinositol 3-kinase (PI3K), p85α and p85β. Oncotarget 8, 55863–55876 (2017).

83. Sato, M., Ueda, Y., Takagi, T. & Umezawa, Y. Production of PtdInsP3 at endomembranes is triggered by receptor endocytosis. Nature Cell Biology 5, 1016–1022 (2003).

84. Alcázar, I. et al. P85β phosphoinositide 3-kinase regulates CD28 coreceptor function. Blood 113, 3198–3208 (2009).

85. Céfaï, D., Schneider, H., Matangkasombut, O., Brody, J. & Rudd, C. E. CD28 Receptor Endocytosis Is Targeted by Mutations That Disrupt Phosphatidylinositol 3-Kinase Binding and Costimulation. 9.

86. Ueki, K. et al. Increased insulin sensitivity in mice lacking p85β subunit of phosphoinositide 3- kinase. Proc. Natl. Acad. Sci. U.S.A. 99, 419–424 (2002).

87. Deane, J. A. et al. Enhanced T Cell Proliferation in Mice Lacking the p85β Subunit of Phosphoinositide 3-Kinase. The Journal of Immunology 172, 6615–6625 (2004).

88. Ueno, T., Falkenburger, B. H., Pohlmeyer, C. & Inoue, T. Triggering Actin Comets Versus Membrane Ruffles: Distinctive Effects of Phosphoinositides on Actin Reorganization. Sci. Signal. 4, (2011).

89. Onuma, H. et al. Rapidly rendering cells phagocytic through a cell surface display technique and concurrent Rac activation. Science Signaling 7, 1–8 (2014).

90. Inoue, T., Heo, W. D., Grimley, J. S., Wandless, T. J. & Meyer, T. An inducible translocation strategy to rapidly activate and inhibit small GTPase signaling pathways. Nature Methods 2, 415–418 (2005).

91. Sato, I. et al. Differential trafficking of Src, Lyn, Yes and Fyn is specified by the state of palmitoylation in the SH4 domain. Journal of Cell Science 122, 965–975 (2009).

92. Zacharias, D. A., Violin, J. D., Newton, A. C. & Tsien, R. Y. Partitioning of Lipid-Modified Monomeric GFPs into Membrane Microdomains of Live Cells. Science 296, 913–916 (2002).

93. Zhou, Y. et al. Lipid-Sorting Specificity Encoded in K-Ras Membrane Anchor Regulates Signal Output. Cell 168, 239–251.e16 (2017).

94. Gradinaru, V. et al. Molecular and Cellular Approaches for Diversifying and Extending Optogenetics. Cell 141, 154–165 (2010).

95. Komatsu, T. et al. Organelle-specific, rapid induction of molecular activities and membrane tethering. Nature Methods 7, 206–208 (2010).

96. Schindelin, J. et al. Fiji: an open-source platform for biological-image analysis. Nature methods 9, 676–682 (2012).

97. Deb Roy, A., et al. Optogenetic activation of Plexin-B1 reveals contact repulsion between osteoclasts and osteoblasts. Nature Communications 8, 15831 (2017).

98. Ershov, D. et al. TrackMate 7: integrating state-of-the-art segmentation algorithms into tracking pipelines. Nat Methods (2022) doi:10.1038/s41592-022-01507-1.

